# More social species live longer, have higher generation times, and longer reproductive windows

**DOI:** 10.1101/2024.01.22.575897

**Authors:** Roberto Salguero-Gómez

## Abstract

The role of sociality in the demography of animals has become an intense focus of research in recent decades. However, efforts to understand the sociality-demography nexus have focused on single species or isolated taxonomic groups. Consequently, we lack generality regarding how sociality associates with demographic traits within the Animal Kingdom. Here, I propose a continuum of sociality, from solitary to tightly social, and test whether this continuum correlates with the key demographic properties of 152 species, from jellyfish to humans. After correction for body mass and phylogenetic relationships, I show that the sociality continuum is associated with key life history traits: more social species live longer, postpone maturity, have greater generation time, and greater probability of achieving reproduction than solitary, gregarious, communal, or colonial species. Contrary to the social buffering hypothesis, sociality does not result in more buffered populations. While more social species have a lower ability to benefit from disturbances, they display greater resistance than more solitary species. Finally, I also show that sociality does not shape reproductive or actuarial senescence rates. This cross-taxonomic examination of sociality across the demography of 13 taxonomic classes highlights keyways in which individual interactions shape most aspects of animal demography.

## Introduction

Quantifying the links between sociality and demography is crucial to understand and forecast the dynamics of animal wildlife populations [1,2]. Indeed, social structures can influence various aspects of an organism’s life, including its survival [3,4] and reproduction [5,6]. As such, the links between sociality and the demography of animal species has gained increasing attention in recent years [7–9], as we strive to unravel the mechanisms underlying population fluctuations and viability [10,11].

Social interactions within a group can significantly influence the key rates that shape organismal fitness: the survival and reproduction of individuals in the group. For instance, cooperative breeding, where individuals assist in the care and rearing of offspring, occurs in many animal species, from meerkats (*Suricata suricatta*; [12] to fairy-wrens (*Malurus cyaneus;* [13]. This behaviour may enhance reproductive success by optimising resource allocation [14] and providing protection against predators [15]. Conversely, in some species, competition for mates within social groups can result in skewed reproductive success [16,17]. Survival patterns within a population can also be intricately linked to sociality. Indeed, social animals often experience reduced predation risk through group vigilance [18] and cooperative defence mechanisms [19]. However, social hierarchies can also increase competition [20], and lower survival [21,22]. As such, sociality has the potential to drastically affect the demography of animal species. Understanding how these contrasting factors interact is essential for predicting and managing population dynamics in socially structured species [23].

Our understanding of how sociality shapes the demography of animals is lacking generality. Specifically, we lack a comprehensive overview of how sociality associates with various demographic attributes of animal populations across a wide range of taxonomic classes - but see [24] for a cross-taxonomic examination of reproductive monopolisation across *ca*. 300 species of wasps, ants, birds, and mammals. Most social comparative studies have focused on birds and mammals [25–28]. In these studies, sociality has been linked to enhanced reproductive success [29], and only weakly with reproductive senescence [27]. Within primates, the impacts of sociality are modulated via complex group hierarchies, where dominance influences access to resources, mating opportunities, and reproductive success [30–32]. Among the carnivora, social species like lions (*Panthera leo*) and spotted hyenas (*Crocuta crocuta*) display complex group structures that influence demographic patterns. In them, cooperative hunting and communal care of young enhance the survival and overall fitness of individuals within the social group [33]. In eusocial insects (*e.g.*, ants, bees, and termites), the within-colony division of labour affects demographic traits via resource allocation, colony defence, and reproduction [34,35]. In certain fish species, such as the clownfish (*Amphiprion* spp.) and cleaner wrasses (*Labroides* spp.), sociality is linked to improved foraging efficiency, reduced predation risk, and increased reproductive success [36].

Sociality is multifaceted. As such, different authors define the degree of sociality depending on the taxonomic group under examination. I argue that the lack of cross-taxonomic categorisation of sociality, together with the historical scarcity of demographic data across the Tree of Life explains why a broader examination of sociality across the demographies of the Animal Kingdom does not exist. However, taking inspiration from the Plant Kingdom, one could think about the individuals in a group of animals as an analogous organisation to the xylem vessels that connect the roots and shoots in an individual vascular plant. The degree of integration-modularity of these individuals (vessels) within the animal group (individual plant) provides a framework to categorise sociality across the Animal Kingdom. Indeed, the degree of integration-modularity across 138 plant species strongly correlates with their longevity and rate of actuarial senescence [37].

Here, I propose a continuum of animal sociality to examine whether sociality, the way individual organisms organise themselves within a population and interact, shapes their demography. My sociality continuum contains five categories: (1) solitary: individuals spend their time alone, except to breed (*e.g.*, tigers (*Panthera tigris*), cheetahs (*Acinonyx jubatus*), some wasps); (2) gregarious: individuals spend time in groups but social interactions are loose (*e.g.*, wildebeests, zebras, flock-forming birds); (3) communal: individuals live together in close proximity and often share a common nesting or dwelling area, but do not engage in cooperative breeding (*e.g.*, purple martin (*Progne subis*)); (4) colonial: individuals live in close proximity and always share a common nesting or living area (*e.g.*, nesting birds, some wasps, coral polyps); and (5) social: individuals live in close proximity and form stable, organised groups, engaging in social behaviours such as cooperative breeding and hierarchical structures (*e.g.*, African elephant (*Loxodonta africana*), most primates, dolphins, meerkat, honeybee (*Apis mellifera*)).

I apply this sociality continuum to 152 species across 13 taxonomic classes, from cnidarians to mammals (Fig. 1), to carry out phylogenetic comparative analyses to examine the demographic correlates of sociality across their life history traits [38], vital rates and impacts on population growth rate [39], and metrics of long-term and short-term performance [40]. My multi-level examination of the demographic traits of these species allows me to test expectations regarding the expected impacts of sociality on longevity and senescence [41] and the ‘social buffer’ hypothesis [42], whereby more social organisms should be able to better mitigate environmental stochasticity while also displaying lower temporal oscillations in population growth rate.

**Figure 1.**
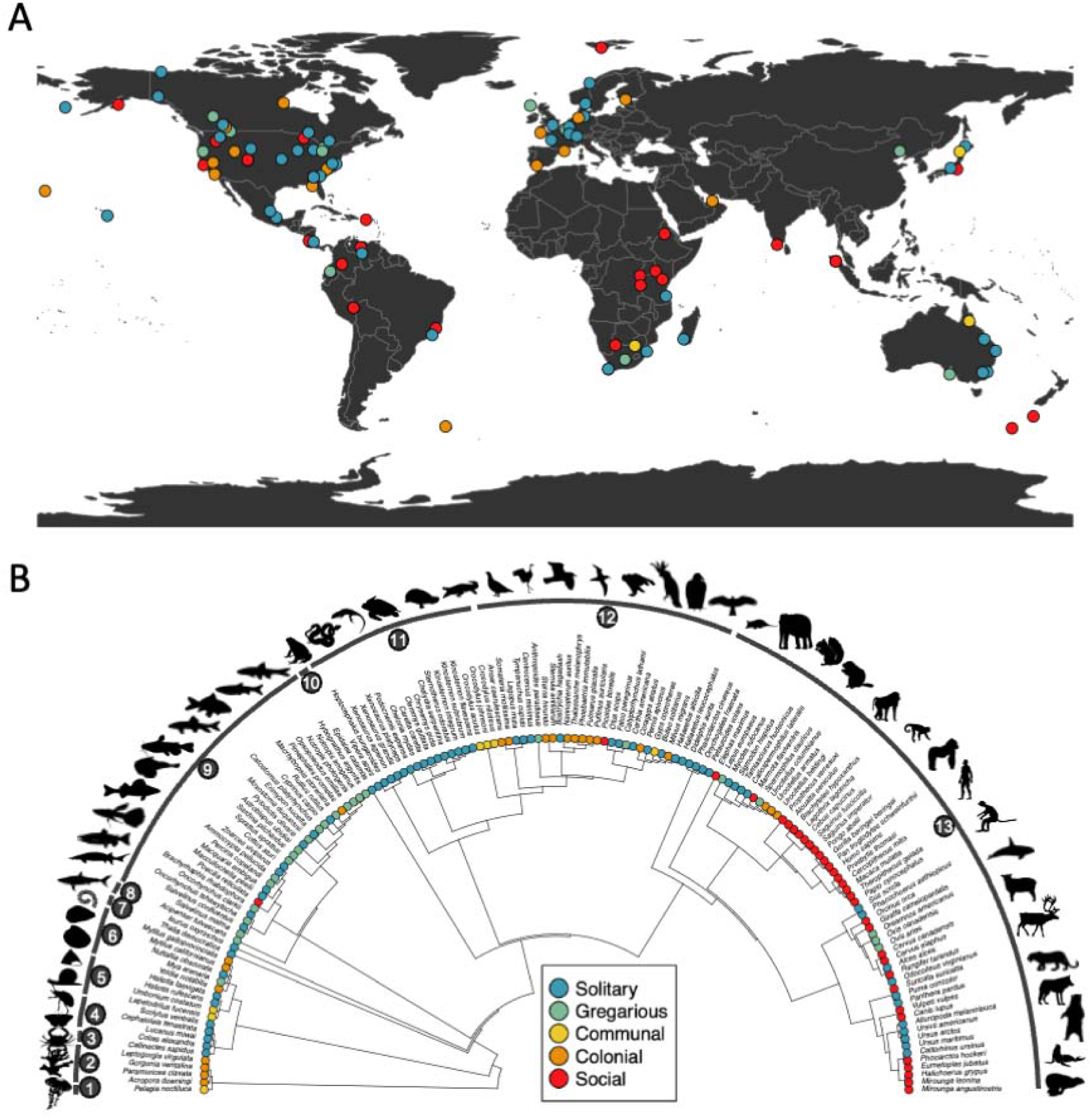
Geographic and phylogenetic representation of the 152 animal species used in this study to examine the correlates of sociality and demographic traits. Species sociality is represented along a discrete continuum, from solitary (blue), gregarious (green), communal (yellow), colonial (orange), to social (red). **A**. World map depicting the geographic location of the examined studies from the COMADRE Animal Matrix Database, when GPS coordinates were available (n = 98 species). **B**. Phylogenetic representation of the 152 study species. Numbered arcs represent the taxonomic classes (from left to right): 1: Scyphozoa (n = 1 species); 2: Anthozoa (n = 4); 3: Malacostraca (n = 1); 4: Insecta (n = 4); 5: Gastropoda (n = 4); 6: Bivalvia (n = 5); 7: Thaliacea (n = 1); 8: Elasmobranchii (n = 1); 9: Actinopterygii (n = 28); 10: Amphibia (n = 1); 11: Reptilia (n = 18); 12: Aves (n = 26); and 13: Mammalia (n = 58). Silhouette source: http://phylopic.org.

## Methods

To test the aforementioned hypotheses, I ran a series of comparative phylogenetic analyses based on high resolution demographic, body mass, sociality, and phylogenetic data of animal species across multiple taxonomic classes.

### Demographic data

To obtain demographic data across a wide range of animal species, I used the COMADRE Animal Matrix Database [43]. COMADRE is an open-access database that contains high-resolution demographic information of animal species, mostly from peer-reviewed publications. In its current version, COMADRE (v. 4.32.1) contains 3,488 matrix population models (MPMs, hereafter [44]) from 429 animal studies across 415 studies. An MPM is a numerical array that describes the demography of a species at a given location, time period, and environmental conditions, in discrete time and along discrete life cycle stages [44]. Specifically, in an MPM, the life cycle of a given population is discretised into stages (*e.g.*, age, discrete size ranges, developmental stages), such that the rates that are then stored in the MPM describe individual-level, stage-specific survival (σ), growth/development (γ), shrinkage/de-development (ρ), and reproduction (φ) from time *t* to *t*+1.

To ensure comparability across the thousands of MPMs available in COMADRE, I imposed the following series of selection criteria using the *Rcompadre* R package [45]: (1) Unmanipulated MPMs from wild populations, so that the derived demographic traits reflect the dynamics of the population under natural conditions, rather than under experimental treatments or in captivity; (2) MPMs with dimension equal to or greater than 4 × 4 (*i.e.*, 4 or more stages in the life cycle), so as to allow for enough resolution in the life cycle to successfully discern age-specific patterns of survivorship and reproduction [46,47] and avoid mortality and fertility plateaus [48]; (3) MPMs where the rates of survival and reproduction had been separated into sub-matrices (such that the overall MPM ***A*** = ***U*** + ***F***, where ***U*** and ***F*** represent the sub-matrices of survival and reproduction, respectively), to facilitate the calculation of life history traits and vital rates and their elasticities to population growth rate; (4) MPMs whose stage-specific survival values ≤ 1, to prevent erroneous calculations of inflated longevity and actuarial senescence [47]; and MPMs corresponding to extant species (*e.g.*, a study successfully passed through selection criteria 1-4, above, but in fact corresponds to an extinct porcupine, *Hystrix refossa* [49]). These criteria resulted in 154 animal species, for which 23 species had at least two available studies in COMADRE.

Next, for those 23 species with multiple studies available, I manually chose the single study per species that provides higher representativity of the demography of the species of interest by retaining the longer-term study, the MPM with greater dimensionality, and/or the study where the rates of survival and/or reproduction in adults were not assumed constant. For each of the resulting 154 animal species, I retained the ‘grand mean’ MPM, which describes the dynamics of the population across the length of the study by calculating the element-by-element arithmetic mean MPM, thus resulting in one MPM per species. In addition, I supplemented my dataset with a recent study on meerkat demography [50], not yet incorporated in COMADRE. For the 10 years of seasonal MPMs available in this study, I retained the MPMs corresponding to 1998-2002, corresponding to normal environmental conditions (M. Paniw, pers. comm.), and then again estimated the element-by-element arithmetic grand mean MPM. This addition increased the count to 155 animal species. The vast majority of MPMs were parameterised with data collected once per year (n = 130, 84%; Fig. S1). However, to make sure that all demographic traits derived from the MPMs (below) were on the same time units, MPMs not on an annual periodicity were rescaled accordingly by elevating each MPM element *a_i,j_*in ***A*** to the power of 1/*P*, where *P* is the projection interval of the MPM, which is also archived in COMADRE [43]. The original sources of the individual demographic studies used in this paper are shown in Table S1.

For each species’ grand MPM, I derived a series of demographic traits that inform the life history traits, vital rates and their relative impacts on population growth rate, as well as metrics of asymptotic (*i.e.*, long-term) and transient (*i.e.*, short-term) performance. To do so, I used the R packages *Rage* [45], *popdemo* [51] and *popbio* [52].

Life history traits are key metrics that define specific moments and periods of time along the life cycle of a species [38]. Previous examinations have demonstrated that the chosen life history traits describe ∼60-80% of the variation in life history strategies of animals [53–57]. The 11 chosen life history traits are:

1. Generation time (*T*) the average age of reproductive individuals in a population [55];
2. Net reproductive output (*R_0_*): the total number of offspring produced by the average individual in the population during its lifespan [44];
3. Mean life expectancy (η*_e_*): mean lifespan of a cohort [44];
4. Variance in life expectancy (Δη*_e_*): variance of lifespan in a cohort [44];
5. Maximum longevity (*L_max_*): time elapsed until 99% of a cohort has died [58];
6. Age at maturity (*L*_α_): age at which the first individual in a cohort reproduces [59];
7. Reproductive window (*L*_α_*_-_*_ω_): the average duration of reproduction of individuals in the population [60];
8. Maturity probability (*p_R_*): the probability of achieving reproduction before dying [44];
9. Actuarial senescence (*s_lx_*): the shape (*s hereafter*) of age-specific survivorship *l_x_* [61,62]; The value of this shape ranges between −0.5 and +0.5: *s_lx_* = 0 indicates negligible actuarial senescence (*i.e.,* constant survival with age), *s_lx_* < 0 indicates negative actuarial senescence (generally increasing survival with age), and *s_lx_*> 0 indicates (positive) actuarial senescence (generally decreasing survival with age);
10. Reproductive senescence (*s_mx_*): the shape of age-specific reproduction *m_x_* [63] describes the symmetry of reproduction over age by comparing the area under a cumulative reproduction curve over age with the area under constant reproduction, and it ranges between −0.5 and +0.5: *s_mx_*= 0 indicates negligible reproductive senescence (*i.e.,* constant reproduction with age), *s_mx_*> 0 indicates (positive) reproductive senescence (decreasing reproduction with age), and *s_mx_* < 0 indicates negative reproductive senescence (increasing reproduction with age);
11. Degree of parity (*S*): quantifies the degree of semelparity (individuals in the population reproduce once; S = 0) *vs.* iteroparity (reproduce multiple times; S>>0) [64] (Fig. S2).

To further characterise the potential associations between sociality and the demography of different animal species, I examined vital rates and their relative impacts on population growth rate. Using the Rage R package [45], I estimated an average value of survival (σ), growth (γ), shrinkage (ρ), and reproduction (φ), weighted by the stationary distribution of individuals in the population [60]. Next, I calculated the elasticities of population growth rate (λ) to these vital rates. These elasticities quantify the effect that a relative, infinitesimally small change in a vital rate would have on the long-term, asymptotics of the population [39,65], quantified as the dominant eigenvector of the MPM ***A***, λ [44] (not to be confused with Pagel’s λ [66], an estimate of phylogenetic signal - below). Next, I also quantified the population growth rate (λ), and the deviance of population growth rate from λ=1 as |1-λ|, at which point the population is neither increasing (λ>1) nor declining (λ>1). Deviance from λ=1 was of particular interest here, as the social buffer hypothesis [42] predicts the canalisation of λ towards demographic stability (λ ≈ 1) with increased sociality.

Finally, I also calculated metrics of transient dynamics. Transient dynamics in an structured population model, such as an MPM, quantify the short-term responses of a population to potential disturbances across the structure of said population [40,67]. The full range of potential responses to disturbances creates a so-called transient envelope [40], from which one can calculate the rate of recovery to the stationary equilibrium (damping ratio, ζ), the number of oscillations that the population undergoes before achieving said equilibrium (period of oscillation, *P_i_*), the maximum increase in size that the population can undertake following a disturbance (reactivity, 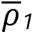), and the maximum decrease in size that the population can display after a disturbance (first step attenuation, 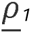). I note that many other transient metrics can be used to define the transient envelope of an MPM [40], but the four I chose here tend to be uncorrelated [68,69], and together they inform the inherent ability of a population to respond to disturbances, that is, their demographic resilience [70].

The vital rate elasticities, and metrics of asymptotic and transient performance were only calculated for MPMs that are reducible, primitive, and ergodic, thus guaranteeing the existence of a single dominant eigenvalue, λ*_1_* [40,44].

### Adult body mass data

Most life history traits scale with adult body mass [53–56]. As such, to explore the relationships between sociality and life history traits beyond the confounding effect of body mass, I first obtained adult body mass data for each species. For mammals, I used information archived in AnimalTraits [71]; for birds, AVONET [72]; for fish, FishBase [73] via rFishBase [74]; for mammals, birds, and reptiles and amphibians, AMNIOTE [75]; and for remaining groups, MOSAIC [76], and data from [54]. For 18 remaining species for which adult body mass data were not available from these online databases (mostly insects, bivalves, and corals), I obtained body mass information from peer-review publications using the keywords “body mass” OR “weight” and the name of the species in ISI WoS. Data were transformed to grams across all sources. The final variable was log-transformed to fulfil assumptions of the statistical analyses employed below.

### Sociality data

I classified sociality following my proposed sociality continuum. In the development of this classification, my goal was to explicitly acknowledge how sociality is not a binary category, and that it indeed has a multi-faceted nature, as animals can vary in their social interactions according to temporal and spatial dimensions [77], as well as a function of group size [78], group hierarchy [79], and type of interactions [80]. My continuum of sociality takes five ordered levels, from less to more social, as follows: (1) Solitary: individuals spend most of their life cycles alone, except to breed; (2) Gregarious: individuals spend some time in groups, but their social interactions are loose, and these interactions can frequently disaggregate; (3) Communal: individuals live in close proximity, often sharing a common nesting or dwelling area, but do not engage in cooperative breeding; (4) Colonial: individuals live in close proximity and always share a common nesting or living area; and (5) Social: individuals live in close proximity and form stable, organised groups, engaging in social behaviours such as cooperative breeding and hierarchical structures. From solitary to social, the degree of spatial and temporal interactions increases, as does the quality of the interaction.

No centralised, open-access database exists to my knowledge offering a comprehensive categorisation of the sociality of species across kingdom Animalia. As such, I classified species according to my continuum of sociality drawing from multiple sources, which allowed me to carefully cross-validate the classification for each species. I first contrasted information from the following online resources for each species in my dataset: Animal Diversity Web [81], Smithsonian’s National Zoo and Conservation Biology Institute (https://nationalzoo.si.edu/), FishBase (https://www.fishbase.se/search.php), IUCN Red List of Threatened Species [82], and chatGPT (https://chat.openai.com/) - for the latter, suggested references were cross-checked independently and validated via ISI WoK. Next, I double-checked existing classifications in my data and complemented missing information from peer-review publications for each of the 154 animal species. Once the list was completed, I discussed it with experts in each of the taxonomic classes, and made the appropriate corrections (Table S1).

### Phylogenetic data

The analyses below explicitly consider how the examined species are related phylogenetically. To that end, I obtained a phylogeny from the Open Tree of Life [83] via the R package *ROTL* [84]. I first matched the scientific names of my subset of animal species from COMADRE to the Taxonomic Name Resolution Service via the function *tnrs_match_names*. All species matched successfully, except the two species from class Demospongiae (*Spongia graminea*, and *Xestospongia muta*). As the phylogenetic tree of Demospongiae is not well resolved [85], I proceeded with the analyses discarding these two species, and so the final analyses include 152 animal species, instead of 154.

### Comparative phylogenetic analyses

To examine the correlates of sociality with the demographic traits examined from the MPMs in COMADRE, I first ran a phyloANOVA via the *phytools* R package [86]. For each demographic trait, the response variable was its residual from the pgls regression [66] over adult body mass, and the explanatory variable was the factor variable sociality continuum, with its five different levels, ordered as “solitary”, “gregarious”, “communal”, “colonial”, and “social”. To correct for type II errors, after the tests were performed, I also implemented a Bonferroni correction with the function *p.adjust* from the *Stats* R package. The phylogenetically-corrected, body mass-corrected post-hoc Tukey test that I implemented next explicitly allowed me to test the hypothesis that the sociality continuum, which ranks species along increasing degree of social interactions, associates with different life history traits, vital rates and their elasticities of population growth rate, as well as metrics of asymptotic and transient dynamics.

The phylogenetic ANOVAs performed here allowed me to examine the correlates of sociality with each life history trait separately. However, life history traits are often correlated with life history strategies [38]. Indeed, previous works have demonstrated that animals show two primary axes of variation: one where individuals either develop fast or live long, the so-called fast-slow continuum [38,53,56,87], and another one along which species differ in their reproductive strategies, the so-called reproductive strategies continuum [10,54,87].

To quantify the life history strategy space of the 152 study animal species, I used a phylogenetic principal component analysis (pPCA). Briefly, PCA is a family of multivariate statistical techniques used to examine complex data by reducing their dimensionality to highlight the main factors that explain observed variation [88]. The pPCA takes into consideration the non-independence of life history traits due to species relatedness [89,90]. To avoid co-linearities in the pPCA, I first examined the pairwise correlations of the 11 life history traits defined above, and chose only one life history trait when the spearman correlation coefficient of a pairwise life history trait correlation was > 0.70. This step led me to exclude from pPCA analyses the variance in life expectancy (Δη*_e_*) and maximum longevity (*L_max_*), which were highly correlated with mean life expectancy (η*_e_*) (0.83 and 0.90, respectively; Fig. S3). Next, because PCA approaches require a dataset with no missing values, and my data contained missing values (Fig. S4), I carried out imputations only on life history traits for which the degree of missing data was <40%. This step further excluded the rates of actuarial (*s_lx_*) and reproductive senescence (*m_lx_*) from the pPCA analyses. These steps identified seven life history traits for the pPCA: T, *R_0_*, η*_e_*, *L*_α_, *L*_α_*_-_*_ω_, *p_R_*, and *S*.

To implement the pPCA, I first log-transformed the aforementioned seven life history traits to fulfil assumptions of error normal distribution necessary in linear models. Next, I calculated the residuals of each life history trait as a function of species’ adult body mass via a pgls with the R package caper [66] and the phylogeny. Next, I rescaled the residuals to mean (μ) = 0 and variance (σ) = 1. This approach allowed to establish primary axes of variation of, in this case, life history traits, as well as to estimate the overall strength of the phylogenetic relationships among species. For the latter, I also estimated Pagel’s λ, a parameter that ranges between 0 (the pattern of association is not explained by the phylogenetic hypothesis), and 1 (the pattern is entirely explained by phylogenetic inertia).

I used a well-established protocol [53,54,60,87] to recover the missing information from the seven life history traits using an iterative PCA approach. Briefly, I employed the “*regularised*” method in the function *mice* in the R package *mice* [91] with 20 sets of imputation. To draw strength across the full dataset, the imputation took into account the vital rate values, elasticities, transient dynamics metrics (described below), and body mass (above) of each species. Here, an iterative algorithm first assigns the mean of each life history trait to the missing values of that life history trait. Next, a regular PCA is performed, and the scores of the species with missing data are re-assigned using the orthogonal relationships of the PCA. This iteration is repeated until convergence. As a check, I (1) re-examined the pairwise correlations of the life history traits pre-(Fig. S3) and post-imputation (Fig. S5), which did not qualitatively change in their covariation structure and (2) implemented the same protocol using only the 85 species for which no life history trait data were missing, and found that the results with imputed data are insensitive to the imputation (not shown). The negligible sensitivity of the results reported here to the imputation approach is in agreement with previous comparative demographic studies that also used COMADRE [10,54,92].

Finally, to examine how the sociality continuum governs the life history strategy space of animals, I used the scores of the pPCA (above) to run further phylogenetically-corrected post-hoc Tukey tests to examine if the different levels of the sociality continuum are significantly clustered along the different PC axes.

## Results

The phylogenetic signal of my sociality continuum, from solitary species, like the giant panda (*Ailuropoda melanoleuca*; Table S1), to social species, like the red howler (*Alouatta seniculus*), across the 152 examined animal species, was high. Indeed the estimates of Pagel’s λ for the sociality continuum across these species is 0.907 (95% C.I.: 0.865-0.945; *P* < 0.001). This strong signal is also visually apparent upon examining the phylogenetic tree shown in Fig. 1B. The phylogenetic signal of the associations between the sociality continuum and the examined demographic traits, in contrast, was variable (Table 1). The estimates of Pagel’s λ from the performed pgls models ranged from virtually absent across some demographic rates (*e.g.*, degree of parity, vital rates of individual growth, individual shrinkage, and reproduction, population growth rate (also called λ [44]), period of oscillation), to high in some life history traits (age at maturity (0.759), reproductive window (0.759), and mean life expectancy (0.745)).

**Table 1.**
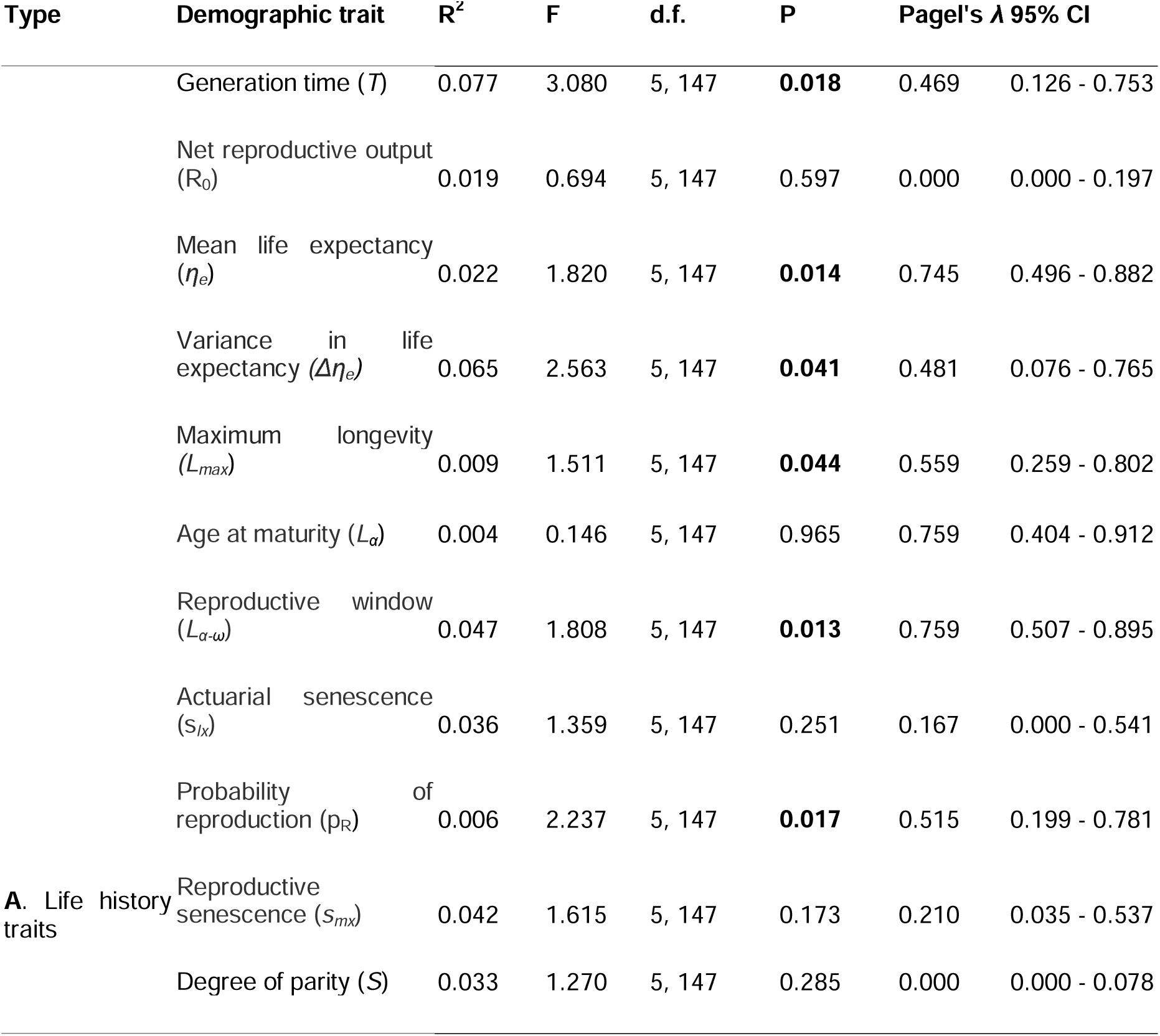
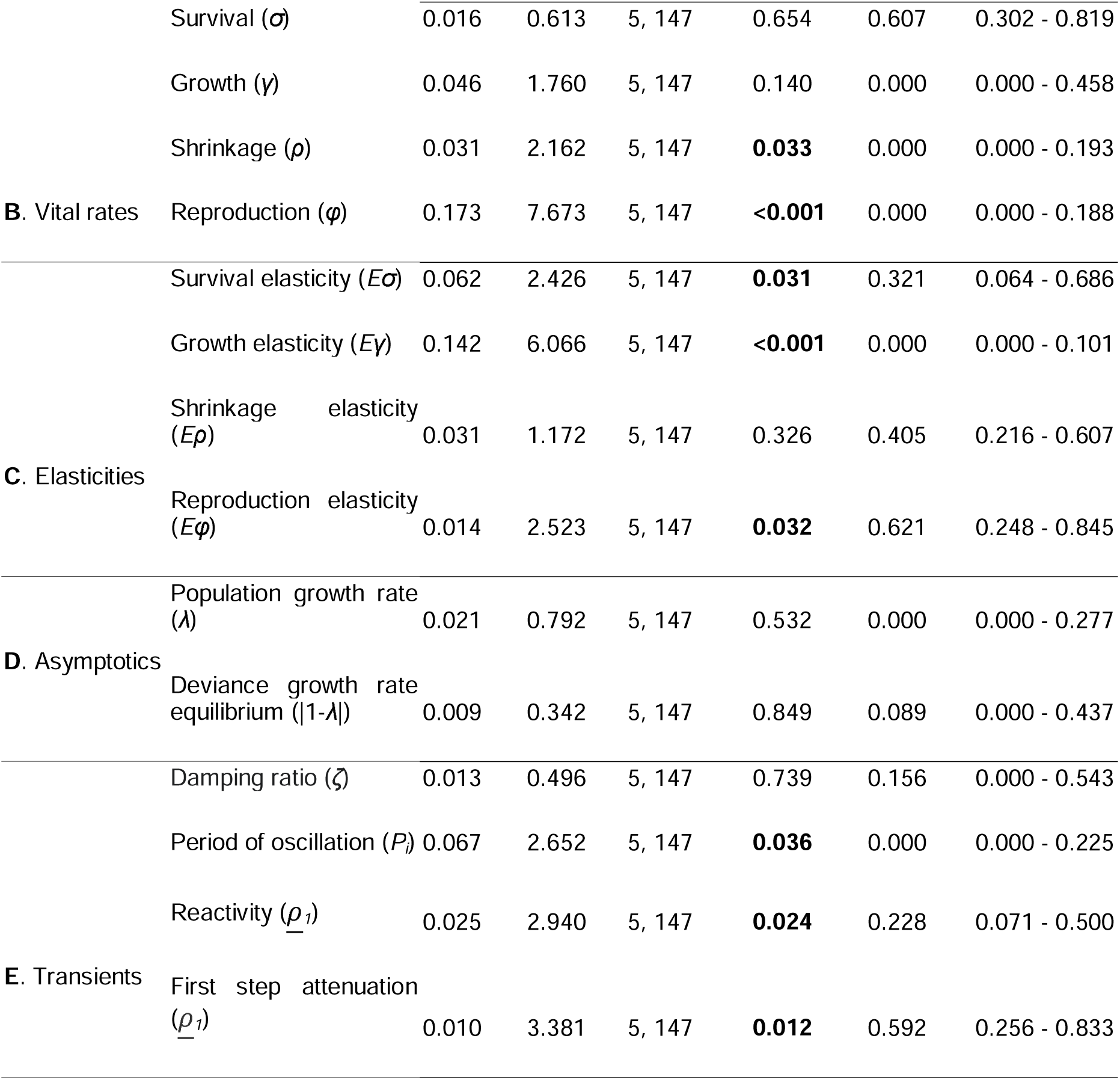
The degree of sociality correlates with key demographic traits across 152 animal species. Phylogenetic generalised least square (pgls) models of sociality and different demographic traits. The demographic traits are classified into five general types: **A**. Life history traits; **B**. Vital rates; **C**. Elasticities of population growth rate (λ) to vital rates; **D**. Metrics of asymptotic (*i.e.,* long-term) performance; and **E**. Metrics transient (*i.e.,* short-term) performance. **Bold**: *P* < 0.05. Pagel’s λ quantifies the phylogenetic signal in the model, and it ranges from 0 (*i.e.,*no phylogenetic signal) to 1 (*i.e.,* pattern completely explained by phylogenetic relationships).

Across the five different types of demographic traits examined here (Table 1: life history traits, vital rates, vital rate elasticity, asymptotics, and transients), the only category where the sociality continuum did not raise some statistically significant results was the metrics of asymptotic dynamics: population growth rate and deviance growth rate equilibrium. Moreover, in these two metrics, the 95% confidence intervals of Pagel’s λ for the respective pgls models overlapped with 0, indicating no phylogenetic effect in the relationship between sociality and long-term population performance. After correcting for adult body mass and phylogenetic relationships, I found that more social species attain longer generation times, longer mean life expectancy, greater variance in life expectancy, higher maximum longevities, and are more likely to reproduce before dying (Table 1A). Interestingly, the relationships between the sociality continuum and species’ vital rates were opposed to their elasticities: more social species have an increased ability to decrease in size and to reproduce (Table 1B), but their population growth rates would increase the most if their vital rates of survival and growth would increase (Table 1C). Finally, for the transient metrics, I found that more social species tend to display greater periods of oscillation, lower reactivity (*i.e.,* propensity to benefit from disturbances, *sensu* [70]), and higher first step attenuation (*i.e.*, resistance to disturbances, *sensu* [70]).

The phylogenetic ANOVA post-hoc Tukey tests revealed important differences in the examined demographic traits as a function of the sociality continuum. For instance, increases in sociality (solitary → gregarious → communal → colonial → social) were associated with prolonged generation time, mean and variance of life expectancy, maximum longevity, longer reproductive windows, and higher probability of reproducing before dying, but not with net reproductive output, age at maturity, actuarial or reproductive senescence, nor degree of parity (Fig. 2). It is worth highlighting that, when the pattern was significant, it did not always change monotonically. Examples include generation time and maximum longevity, where solitary species attained the same range of values as social species, whereas the next level on sociality, gregarious species, had significantly lower values than those of more social species (Fig. 2). Social species also had higher survival as well as lower individual-level shrinkage and reproduction compared to most other categories (Fig. 3). These patterns remained largely consistent when evaluating the elasticities of population growth rate to said vital rates. Finally, for the transient metrics, overall, as the degree of sociality increased, so did the period of oscillation and first step attenuation, but species’ reactivity decreased (Fig. 4). Here, again, we note that the range of values of solitary species do not differ from social species for the period of oscillation and reactivity, but do differ between the next category, gregarious, and the next three categories: communal, colonial, and social.

**Figure 2.**
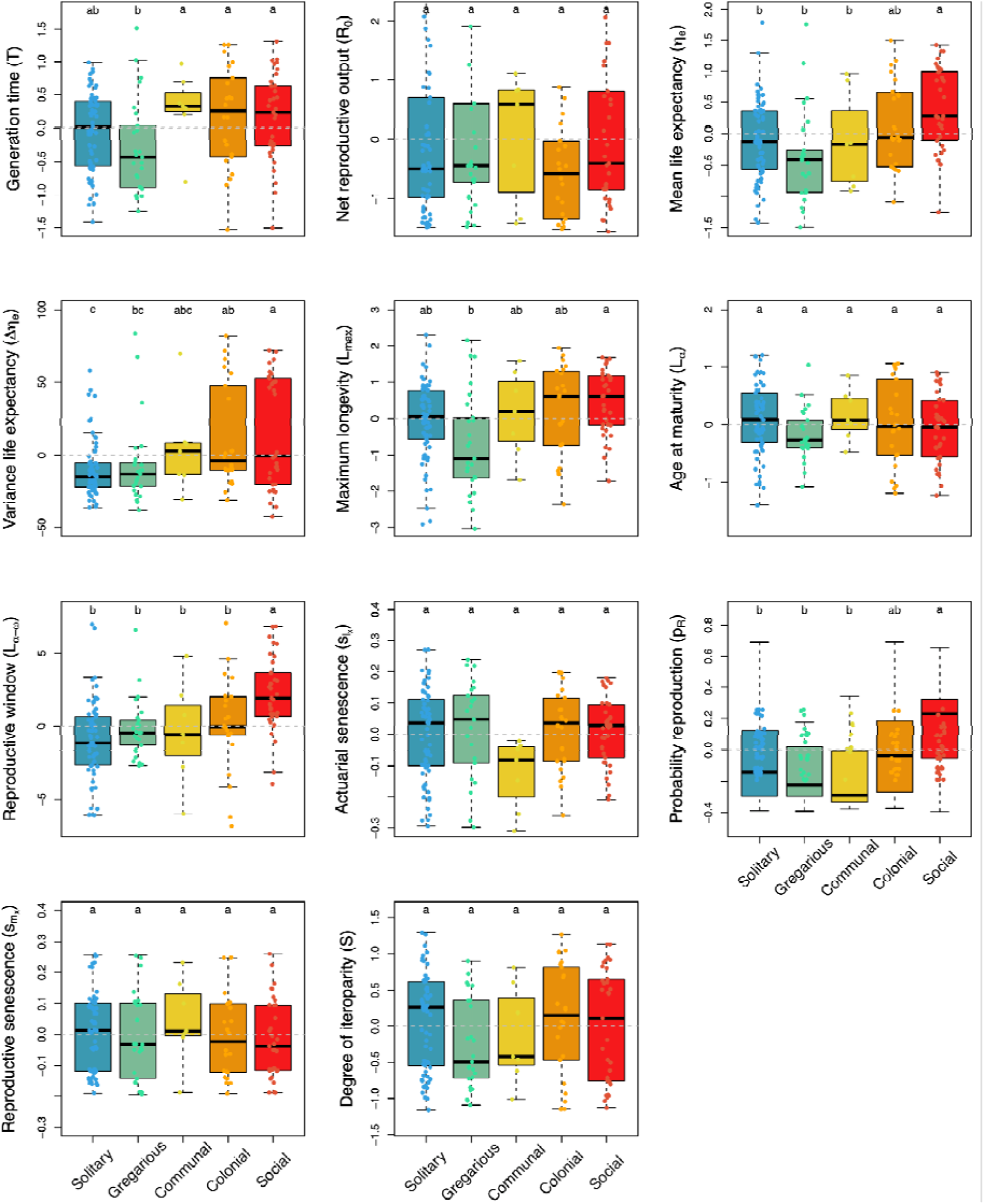
The continuum of sociality correlates with key life history traits. Boxplots of the residuals of life history traits after correction for adult body mass and phylogenetic relationships as a function of the discrete continuum of sociality for 152 animal species. The letters above each group represent post-hoc Tukey test significance levels from the phylogenetic ANOVA. Groups with different letters are significantly different at *P* < 0.05. The horizontal grey dashed line indicates no effect. See Table 1 for full statistics.

**Figure 3.**
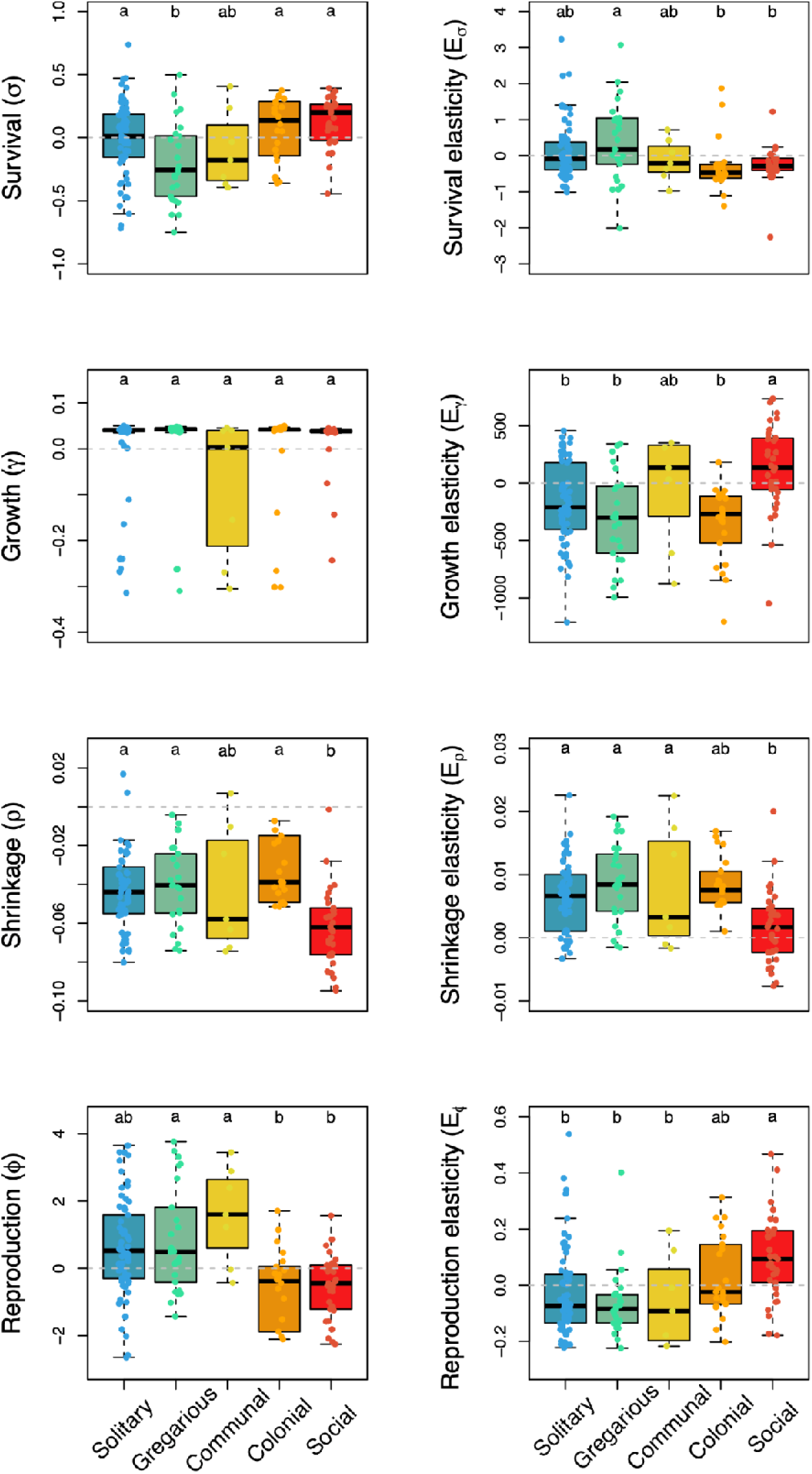
The continuum of sociality correlates with key vital rates and their elasticities to population growth rate. Boxplots of the residuals of vital rates (left: survival σ, growth γ, shrinkage ρ, and reproduction φ), and their elasticities to population growth rate λ (right) after correcting for adult body mass and phylogenetic relationships, as a function of the discrete continuum of sociality for 152 animal species. The letters above each group represent post-hoc Tukey test significance levels from the phylogenetic ANOVA. Groups with different letters are significantly different at *P* < 0.05. The horizontal grey dashed line indicates no effect. See Table 1 for full statistics.

**Figure 4.**
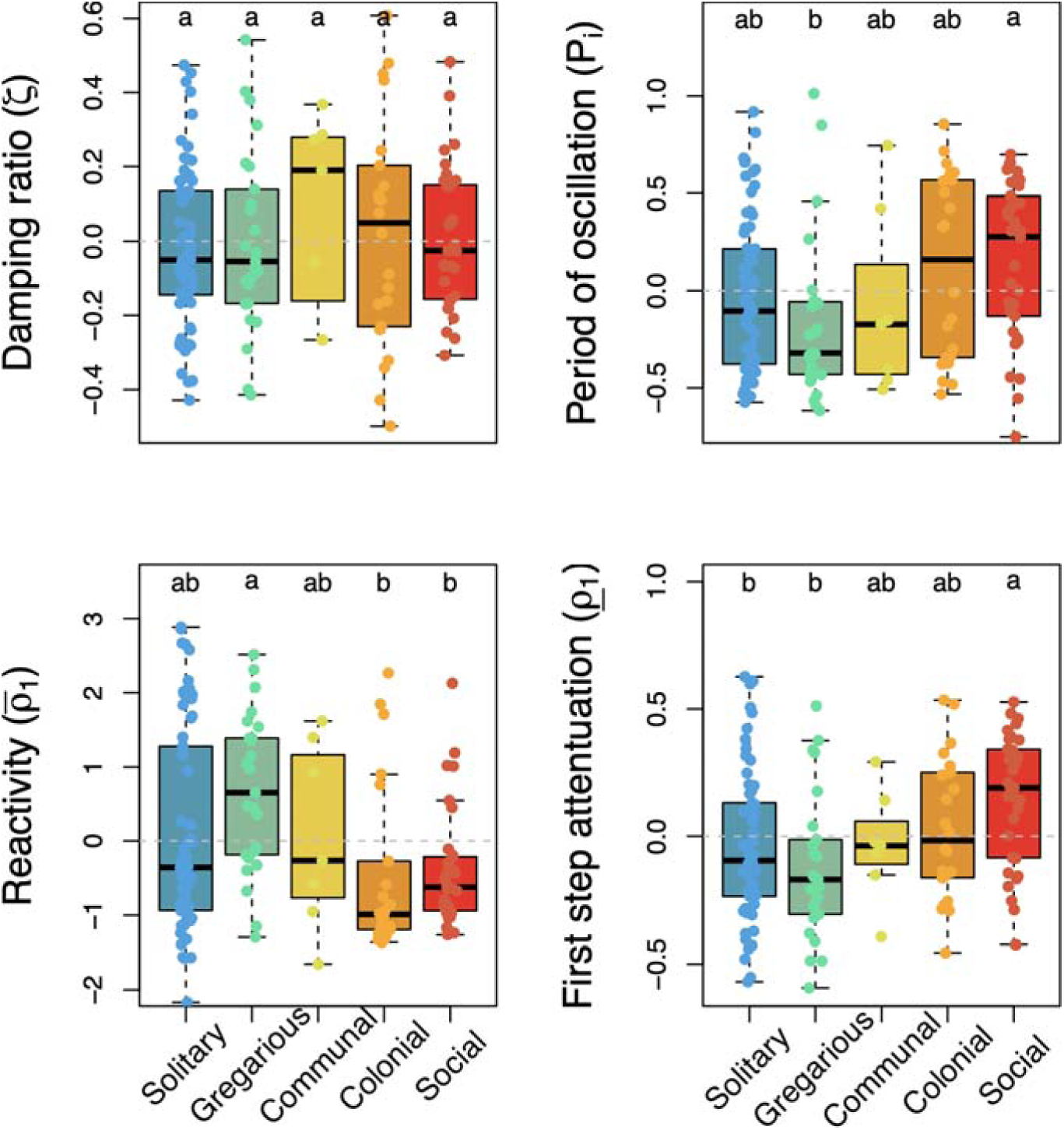
The continuum of sociality correlates with key attributes of transient (short-term) dynamics that predict the inherent ability of natural populations to respond to disturbances. Boxplots of the residuals of four transient dynamics metrics after correction for adult body mass and phylogenetic relationships, as a function of the discrete continuum of sociality for 152 animal species. Damping ratio (ζ) quantifies the speed of recovery post disturbance; Period of oscillation (*P_i_*) quantifies the number of cycles a population goes through before achieving asymptotic dynamics; Reactivity (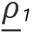) quantifies how much a population can grow after one time step following a disturbance, while first step attenuation (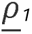) quantifies the loss of resistance after one time step. The letters above each group represent post-hoc Tukey test significance levels from the phylogenetic ANOVA. Groups with different letters are significantly different at *P* < 0.05. The horizontal grey dashed line indicates no effect. See Table 1 for full statistics.

The life history strategy space defined by the seven examined life history traits (Table 1A) identified two principal components whose associated eigenvalues > 1. Following Kaiser’s criterion [93], I retained these two axes for the next part of the analysis. PC1 explains 33.72% of the variation in life history traits across the 152 animal species, and it separates species with low (left of Fig. 5) *vs.* high reproductive schedules. Indeed, species to the right in the pPCA attain higher net reproductive outputs (*R*_0_), have longer reproductive windows (*L*_α_*_-_*_ω_), higher probability of reproducing before dying (*p_R_*), and a higher degree of iteroparity (*S*). PC2 explains 26.20% of the variation, and separates species that live long *vs.* short. Along PC2, species at the top postpone reproduction (*L*_α_), have greater generation times (T), and live longer (η*_e_*) than those at the bottom. The emergent phylogenetic signal, after adult body mass correction, in this phylogenetic PCA was relatively low, with an estimated Pagel’s λ of 0.420 (0.341-0.689 95% C.I.).

**Figure 5.**
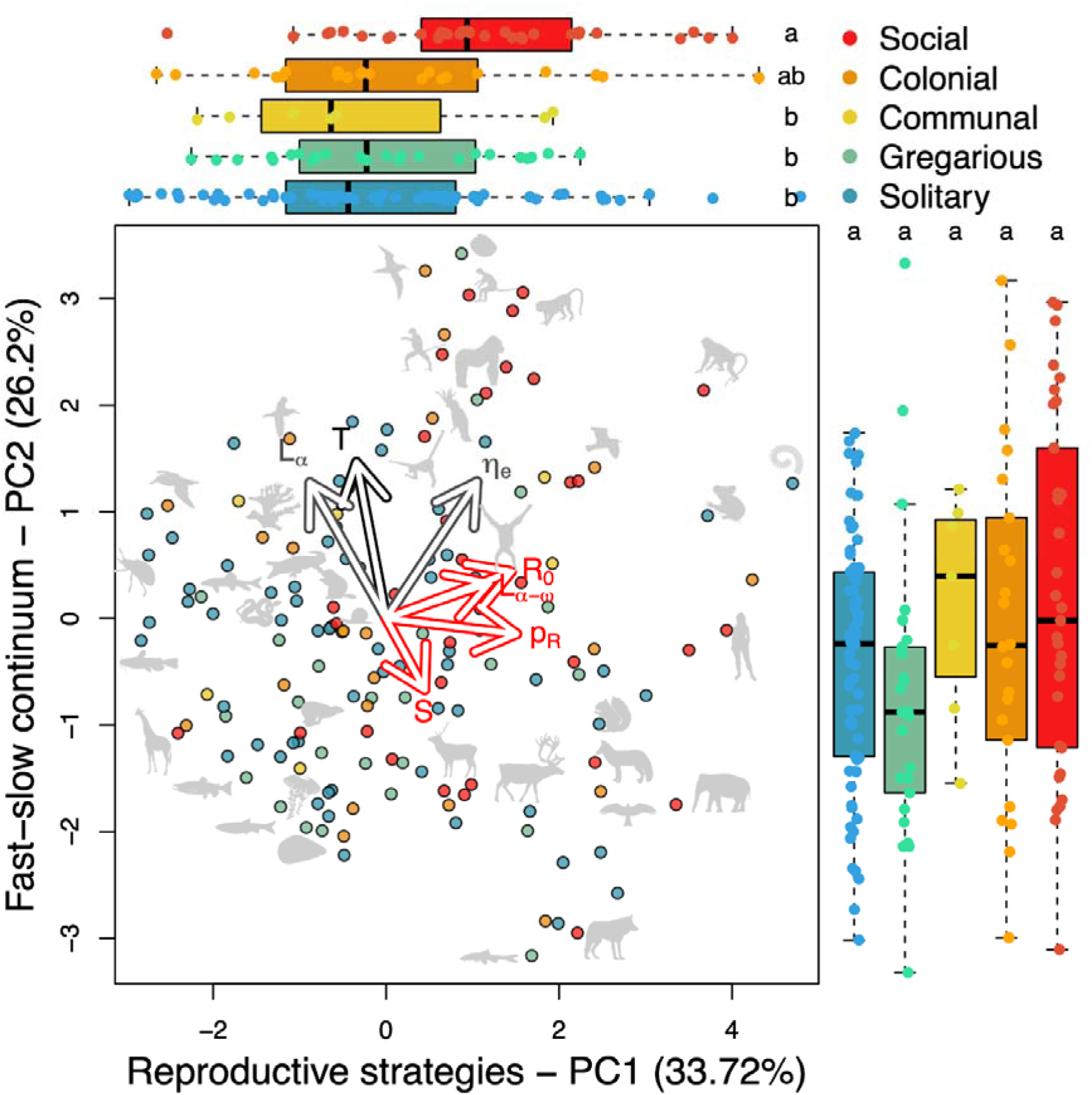
The life history strategy space of animals predicts the placing of sociality along the reproductive strategies continuum, but not the fast-slow continuum. Adult body-mass, phylogenetically corrected principal component analyses (pPCA) of seven life history traits shown in Table 1A and Figure 2. These traits are colour-coded by investments in survival and reproduction (black - generation time, *T*), survival only (grey - age at maturity, *L*_α_; mean life expectancy, η*_e_*, or reproduction only (red - net reproductive output, *R_0_*, reproductive window, *L*_α_*_-_*_ω_; probability of achieving reproduction before dying, *p_R_*; degree of iteroparity, *S*). Each point in the PCA space corresponds to an animal species, with some represented by a silhouette (source: http://phylopic.org; refer to Fig. 1B). The loadings of the life history traits identifies: PC1 (explaining 33.72% of the variance) as the reproductive strategies continuum, with species with grater net reproductive output (*R_0_*), longer reproductive windows (*L*_α_*_-_*_ω_), higher probability to reproducing before dying (*p_R_*), and higher degree of iteroparity (*S*) to the right; and PC2 (explaining 26.20% of the variance) as the fast-slow continuum, with species living longer (η*_e_*), postponing their first reproductive event (*L*_α_), and having greater generation times (*T*) towards the top. The side boxplots correspond to the groups along PC1 (top) and PC2 (right) as a function of the discrete continuum of sociality.

The ranking of species along the sociality continuum raised detectable differences along PC1 (the reproductive strategies continuum; *P* = 0.001; Fig. 5). Indeed, social species like the macaque (*Macaca mulatta*) or the Asian elephant (*Elephas maximus*) or humans (*Homo sapiens*), and colonial species like the common tern (*Sterna hirundo*) are associated with longer reproductive life histories, compared to solitary species like the flathead catfish (*Pylodictis olivaris*) or the common snapping turtle (*Chelydra serpentina*). I found no association between the sociality continuum and the fast-slow continuum, PC2 (*P* = 0.107).

## Discussion

Sociality, the degree to which animals engage in interactions and form cohesive structures, can shape their demography (*i.e.* dynamics of population size and structure) via impacts on their rates of survival [94] and reproduction [16], the building blocks of fitness. Animals show a wide range of social behaviours, including cooperative breeding [13], group living [3], and hierarchical structures that may better protect their young [32], their territory [95], *etc*. Though much research has examined the demographic correlates of animal sociality, these investigations are typically constrained by taxonomic siloes [96]. Indeed, much animal sociality research focuses on single species, or on mammals [4] or birds [27]. Here, by proposing a sociality continuum that ranks species according to the type, spatial, and temporal extent of their interactions, from solitary, to gregarious, to communal, to colonial, to social, I show that, across 152 animal species from 13 taxonomic classes, as the degree of these interactions increases, species life histories attain longer, more iteroparous windows of reproduction[38], even after accounting for the allometric effects of body mass [54,68] and potential constraints of phylogenetic inertia [89,90]. My comparative phylogenetic analyses also show that these social interactions do not shape long-term metrics of performance, such as population growth rate, contrary to the ‘social buffer’ hypothesis [42], but that the sociality continuum is associated with some key attributes of demographic resilience [67].

A wealth of scientific evidence highlights the profound impacts of sociality on life history traits in animals. Indeed, social structures often influence reproductive strategies, with cooperative breeding systems, for example, promoting delayed sexual maturity and extended parental care [97]. In species exhibiting eusociality, such as certain bees and ants, caste systems shape life history trajectories for individuals within the colony [98,99]. Sociality can also affect reproductive success, as observed in the intricate mating systems of various primates [100] and birds [13,101]. Moreover, the presence of social alliances may impact longevity and survivorship, providing advantages in terms of predation avoidance and access to resources [32,80]. Here, I show that the sociality continuum is associated with key life history traits such as prolonged generation time, postponed age at maturity, and increased mean, maximum, and variance of longevity, longer reproductive windows, and greater probability of achieving reproduction before dying. It is worth noting, however, that the interplay between sociality and life history traits extends beyond reproduction and survival, influencing factors like dispersal patterns [102] not taken into account in the present study. To understand this complexity, future work must consider species-specific adaptations and ecological contexts and their dispersal/migration abilities, employing a combination of field observations, experimental manipulations [103], and modelling approaches based on big data [104].

In the present study, the sociality continuum does not correlate with the rate of actuarial nor reproductive senescence across the examined animal species. The impacts of sociality on senescence in animals has been a subject of extensive research, with findings offering mixed perspectives [3]. Studies on various species indicate that social interactions can either accelerate [105] or decelerate senescence processes [106]. For instance, in certain cooperative breeding systems, where individuals collectively care for offspring, sociality may delay reproductive senescence due to shared parental responsibilities and increased chances of offspring survival [27,107,108]. Conversely, in some species with intense intrasexual competition, the pressure to secure mates and resources may accelerate reproductive senescence [109]. Actuarial senescence, reflected in declining survival rates with age, can also be influenced by social factors. For example, the presence of social allies may enhance protection against predation, potentially extending lifespan [80]. However, the stress associated with social hierarchies and conflicts can have the opposite effect, accelerating actuarial senescence [110]. The complex interplay between social dynamics and senescence, and the multiple ways through which senescence can manifest itself [111], underscore the need for a nuanced understanding that considers species-specific traits and environmental contexts. Ultimately, a combination of the comparative approach used here (to identify patterns and plausible proximal mechanisms), coupled with experimental manipulations (to identify and understand distal mechanisms), will provide the ultimate understanding of the conditions under which sociality accelerates or slows down senescence in animals.

Similar approaches will be necessary to disentangle the direction of causality between sociality and demography. The macroecological approach presented here identifies patterns and likely candidate species for further study (*e.g.*, species around the periphery of the life history strategy space presented in Fig. 5) but, as any macroecological exercise [112], mine cannot pinpoint the exact mechanism nor imply causality in the relationships I report. Nonetheless, some comparative work in birds has shown that families with high proportions of cooperation have high survival values, even in non-cooperative breeders in the family [13], suggesting a demography → sociality directionality. Indeed, phylogenetic approaches exist to reconstruct ancestral trait values along phylogenetic hypotheses [113], which would allow one to explore whether shifts in life history traits cause or are followed by changes in sociality along species lineages [114]. However, I argue that these approaches bring too much uncertainty, as both life history traits and sociality are too labile for phylogenetic reconstructions to produce accurate estimates [89]. Instead, I propose that the best way to prove causality between degree of sociality and species’ demographies requires the collective effort of behavioural biologists, examining populations under the same set of control and treatment conditions, and to follow multiple generations.

The relationship between sociality and the ability of animals to withstand environmental stochasticity has garnered substantial attention. Numerous studies suggest that sociality can serve as a buffering mechanism [42], providing a range of benefits that enhance fitness in stochastic environments. While the social buffer hypothesis predicts that sociality should confer adaptive advantages in mitigating the impact of environmental stochasticity, I found that sociality is not associated with any of the metrics of long-term population performance that are often used as population-level proxies for fitness [44,115]. For instance, when evaluating how far from a population growth rate at equilibrium the 152 species’ population were, there was no detectable correlation with sociality. I did, however, find that while more social species do not take a different amount of time to recover after a disturbance compared to less social species, the period of oscillation back to stationary dynamics for the former is greater than in less social species, implying a potentially greater risk of extinction for social species [116,117]. This finding is at odds with the suggestion that sociality increases plasticity against climatic oscillations [118]. Further work integrating phenotypic plasticity and demographic schedules should be key in this context [119]. In this context too, future work should examine how the degree of sociality may vary across populations within the same species. Likewise, the reactivity of social species was lower than in gregarious species, suggesting that, when a population is hit by a disturbance, it is less likely to take advantage by increasing its population size [70]. The past decades have witnessed the development of indexes of demographic buffering based on structured population models [120–123]. Future research should evaluate whether the sociality continuum is intrinsically linked to a potential demographic lability-buffering continuum [124].

Species are more than just social or not social. Thus, a continuum to classify animal sociality is of high appeal. A previous attempt has, however, highlighted some key challenges [24]. Indeed, sociality is shaped by multiple axes, including group size, the kind, timing, and hierarchical nature of interactions, or the partition of labour, to mention a few. The inspiration for my proposed sociality continuum comes from work my laboratory and I carried out to examine the anatomical correlates of actuarial senescence [37]. Finch [125] hypothesised that the ability to compartmentalise risk may allow species to postpone the onset of senescence. Classifying the compartmentalisation of 138 plants (proxied by the distance between their xylem vessels) and 151 animals (proxied by the degree of open-closed circulatory and respiratory systems, and degree of organ redundancy), we found that modularity does predict actuarial senescence in plants but not in animals. Our work there was only focused on vascular plants, for which a standard measure of xylem distance can be obtained [126]. The classification of a continuum of modularity-integration in animals proved difficult then because of the wide range of variation in animal anatomy. Another way to think about animal anatomy is to examine the ‘anatomy’ of their groups and populations: how close are their individuals, how often do they come in contact, and what kinds of interactions do they have. In this regard, Albery and colleagues recently found that older, more senescent (*i.e.*, likely to die) individuals of red deer (*Cervus elaphus*) become more isolated in the group [8], perhaps as a way to contain risk too, as plants do [37]. My proposed sociality continuum correlates with key aspects of the demography of animals. However, further work should examine orthogonal ways in which animal societies evolve, as the predictive power of the correlations examined here remains low (Table 1).

The future of research on animal sociality and demography should prioritise interdisciplinary approaches to analyse a large number of longitudinal studies. This approach will help us unravel the nuanced complexities of social structures and their demographic implications. Integrating advanced technologies, such as GPS tracking [127], social network analysis [8], and comparative genomics [128], can provide a more comprehensive understanding of the dynamic interactions within animal societies. Embracing comparative analyses across a diverse array of taxa can uncover general principles while acknowledging species-specific nuances. As conservation issues become increasingly urgent, linking social and demographic research to conservation biology can inform strategies for preserving biodiversity in the face of environmental challenges [129]. This multidimensional approach holds the key to advancing our understanding of the intricate interplay between sociality and demography in the animal kingdom.

## Supporting information

SOM

## Acknowledgments

I thank J. Firth and K. Davis for conversations regarding the interface of animal sociality in demography; J. Cant and B. Sheldon for input into the classification of sociality of fishes and birds, respectively; the thousands of researchers who have contributed open-access data to the COMADRE Animal Matrix Database (www.compadre-db.org); M. Paniw and E. Conquet for providing access to the demographic models of *Suricata suricatta*; K. Davis for orthographic edits; and G. Albert, J-M. Gaillard, and two anonymous reviewers for constructive feedback on an earlier version of this work I was supported by a NERC Pushing the Frontiers grant (NE/X013766/1).

