## Supplementary material for "More social species live longer, have higher generation times, and longer reproductive windows": SOM

Roberto Salguero-Gómez

**Table S1.** Information about the original source from where demographic information was obtained via the COMADRE database, as well as categorisation of the sociality continuum, and adult body mass (in log, grams).

| **Latin name** | **Common name** | **Class** | **Family** | **Authors** | **Journal** | **Year of publication** | **DOI/ISBN** | **Sociality** | **Adult body mass** |
| --- | --- | --- | --- | --- | --- | --- | --- | --- | --- |
| *Acipenser fulvescens* | Lake sturgeon | Actinopterygii | Acipenseridae | Vélez-Espino; Koops | N Am J Fish Manage | 2009 | 10.1577/M08-034.1 | Gregarious | 11.736 |
| *Acropora downingi* | NA | Anthozoa | Acroporidae | Riegl; Purkis | Glob Change Biol | 2015 | 10.1111/gcb.13014 | Colonial | 10.309 |
| *Ailuropoda melanoleuca* | Giant panda | Mammalia | Ursidae | Carter; Ackleh; Leonard; Wang | Ecol Model | 1999 | 10.1016/S0304-3800(99)00145-3 | Solitary | 11.674 |
| *Alces alces* | Moose | Mammalia | Cervidae | Kvalnes; Saether; Haanes; Røed; Engen; Solberg | Evolution | 2016 | 10.1111/evo.12952 | Solitary | 12.769 |
| *Alouatta seniculus* | Red howler | Mammalia | Atelidae | Wiederholt; Fernandez-Duque; Diefenbach; Rudran | Ecol Model | 2010 | 10.1016/j.ecolmodel.2010.06.026 | Social | 8.764 |
| *Ammocrypta pellucida* | Eastern sand darter | Actinopterygii | Percidae | Finch; Vélez-Espino; Doka; Power; Koops | Fish & Oceans Can | 2011 | NA | Gregarious | -0.223 |
| *Anthropoides paradiseus* | Blue crane | Aves | Gruidae | Altwegg; Anderson | Funct Ecol | 2009 | 10.1111/j.1365-2435.2009.01563.x | Gregarious | 8.537 |
| *Astroblepus ubidiai* | Andean catfish | Actinopterygii | Astroblepidae | Vélez-Espino | Ecol Freshw Fish | 2005 | 10.1111/j.1600-0633.2005.00084.x | Gregarious | 2.688 |
| *Bostrychia hagedash* | Hadeda ibis | Aves | Threskiornithidae | Duckworth; Altwegg; Harebottle | J Ornithol | 2011 | 10.1007/s10336-011-0758-2 | Solitary | 7.127 |
| *Brachyrhaphis rhabdophora* | Live-bearing fish | Actinopterygii | Poeciliidae | Johnson; Zúñiga-Vega | Ecology | 2009 | 10.1890/07-1672.1 | Gregarious | 2.708 |
| *Brachyteles hypoxanthus* | Northern muriqui | Mammalia | Atelidae | Morris; Altmann; Brockman; Cords; Fedigan; Pusey; Stoinski; Bronikowski; Alberts; Strier | Am Nat | 2011 | 10.1086/657443 | Social | 9.291 |
| *Buteo solitarius* | Hawai'ian hawk | Aves | Accipitridae | Klavitter; Marzluff; Vekasy | J Wildlife Manage | 2003 | 10.2307/3803072 | Solitary | 6.225 |
| *Callinectes sapidus* | Blue crab | Malacostraca | Portunidae | Miller | Estuaries | 2001 | 10.2307/1353238 | Solitary | 6.397 |
| *Callorhinus ursinus* | Northern fur seal | Mammalia | Otariidae | Barlow; Boneng | Mar Mammal Sci | 1991 | 10.1111/j.1748-7692.1991.tb00550.x | Solitary | 10.924 |
| *Callospermophilus lateralis* | Golden-mantled ground squirrel | Mammalia | Sciuridae | Hostetler; Kneip; Van Vuren; Oli | PLOS ONE | 2012 | 10.1371/jourNAl.pone.0034379 | Solitary | 5.426 |
| *Calyptorhynchus lathami* | Glossy Black Cockatoo | Aves | Psittacidae | Harris; Fordham; Mooney; Pedler; Araújo; Paton; Stead; Watts; Akçakaya; Brook | J Appl Ecol | 2012 | 10.1111/j.1365-2664.2012.02163.x | Gregarious | 6.138 |
| *Canis lupus* | Gray wolf | Mammalia | Canidae | Miller; Jensen; Hammill | Ecol Model | 2002 | 10.1016/S0304-3800(01)00493-8 | Social | 10.029 |
| *Caretta caretta* | Loggerhead turtle | Reptilia | Cheloniidae | Warden; Haas; Richards; Rose | Ecol Model | 2015 | 10.1016/j.ecolmodel.2014.11.025 | Solitary | 11.600 |
| *Catostomus platyrhynchus* | Mountain sucker | Actinopterygii | Catostomidae | Young; Koops | Fish & Oceans Can | 2013e | NA | Gregarious | 5.521 |
| *Cebus capucinus* | White-faced capuchin monkey | Mammalia | Cebidae | Morris; Altmann; Brockman; Cords; Fedigan; Pusey; Stoinski; Bronikowski; Alberts; Strier | Am Nat | 2011 | 10.1086/657443 | Social | 7.560 |
| *Centrocercus minimus* | Gunnison sage-grouse | Aves | Stercorariidae | Davis; Hooten; Phillips; Doherty | Ecol Evol | 2014 | 10.1002/ece3.1290 | Solitary | 7.409 |
| *Cephaloleia fenestrata* | Neotropical beetle | Insecta | Chrysomelidae | Johnson; Horvitz | Ecology | 2005 | 10.1890/04-0974 | Solitary | 2.485 |
| *Cercopithecus mitis* | Blue monkey | Mammalia | Cercopithecidae | Morris; Altmann; Brockman; Cords; Fedigan; Pusey; Stoinski; Bronikowski; Alberts; Strier | Am Nat | 2011 | 10.1086/657443 | Social | 7.107 |
| *Certhia americana* | Brown Creeper | Aves | Certhiidae | Wintle; Bekessy; Venier; Pearce; Chisholm | Conserv Biol | 2005 | 10.1111/j.1523-1739.2005.00276.x | Solitary | 2.011 |
| *Cervus canadensis* | Rocky Mountain elk | Mammalia | Cervidae | Clark | Thesis | 2014 | NA | Social | 12.206 |
| *Cervus elaphus* | Red deer | Mammalia | Cervidae | Raithel; Kauffman; Pletscher | J Wildlife Manage | 2007 | 10.2193/2005-608 | Social | 11.984 |
| *Chelonia mydas* | Green sea turtle | Reptilia | Cheloniidae | Chaloupka | Book Chapter | 2004 | 978-0-19-803726-2 | Solitary | 11.128 |
| *Chelydra serpentina* | Common snapping turtle | Reptilia | Chelydridae | Congdon; Dunham; van Loben Sels | Am Zool | 1994 | 10.1093/icb/34.3.397 | Solitary | 8.542 |
| *Anser caerulescens* | Snow goose | Aves | Anatidae | Cooch; Rockwell; Brault | Ecol Monogr | 2001 | 10.1890/0012-9615(2001)071[0377:RAODRT]2.0.CO;2 | Colonial | 7.875 |
| *Chrysemys picta* | Painted turtle | Reptilia | Emydidae | Tinkle; Congdon; Rosen | Ecology | 1981 | 10.2307/1941498 | Solitary | 5.918 |
| *Clemmys guttata* | Spotted turtle | Reptilia | Emydidae | Feng; Ross; Mauger; Dreslik | Diversity | 2019 | 10.3390/d11120226 | Solitary | 7.679 |
| *Myodes rufocanus* | Gray-sided vole | Mammalia | Muridae | Yoccoz; Nakata; Stenseth; Saitoh | Res Popul Ecol | 1998 | 10.1007/BF02765226 | Solitary | 3.912 |
| *Colias alexandra* | Queen Alexandra's sulphur | Insecta | Pieridae | Hayes | Oecologia | 1981 | 10.1007/BF00349187 | Solitary | 1.386 |
| *Coragyps atratus* | Black vulture | Aves | Cathartidae | Blackwell; Avery; Watts; Lowney | J Wildlife Manage | 2007 | 10.2193/2006-146 | Colonial | 7.640 |
| *Cottus aturi* | Rocky Mountain Sculpin | Actinopterygii | Cyprinidae | Young; Koops | Fish & Oceans Can | 2013e | NA | Solitary | 4.828 |
| *Crocodylus acutus* | American crocodile | Reptilia | Crocodylidae | Richards | Biological Sciences | 2003 | NA | Communal | 11.806 |
| *Crocodylus johnsoni* | Freshwater crocodile | Reptilia | Crocodylidae | Tucker | Thesis | 1997 | NA | Communal | 9.878 |
| *Crocodylus niloticus* | Nile crocodile | Reptilia | Crocodylidae | Hutton | PhD thesis | 1984 | NA | Communal | 11.453 |
| *Cyprinus carpio* | Common carp | Actinopterygii | Cyprinidae | Stratford; Pollino; Brown | Environ Modell Softw | 2016 | 10.1016/j.envsoft.2016.02.009 | Gregarious | 6.296 |
| *Didelphis aurita* | Common marsupial | Mammalia | Didelphidae | Ferreira; Kajin; Vieira; Zangrandi; Cerqueira; Gentile | Mamm Biol | 2013 | 10.1016/j.mambio.2013.03.002 | Solitary | 7.047 |
| *Elephas maximus* | Asian elephant | Mammalia | Elephantidae | Goswami; Vasudev; Oli | Biol Conserv | 2014 | 10.1016/j.biocon.2014.05.026 | Social | 14.985 |
| *Epidalea calamita* | Natterjack toad | Amphibia | Bufonidae | Di Minin; Griffiths | Ecography | 2011 | 10.1111/j.1600-0587.2010.06263.x | Solitary | 2.944 |
| *Erimyzon sucetta* | Lake chubsucker | Actinopterygii | Catostomidae | Young; Koops | Fish & Oceans Can | 2011 | NA | Solitary | 10.404 |
| *Eumetopias jubatus* | Northern sea lion; Steller sea lion | Mammalia | Otariidae | Holmes; York | Conserv Biol | 2003 | 10.1111/j.1523-1739.2003.00191.x | Social | 12.854 |
| *Falco peregrinus* | Peregrine falcon | Aves | Falconidae | Altwegg; Jenkins; Abadi | Ibis | 2013 | 10.1111/ibi.12125 | Solitary | 6.572 |
| *Fulmarus glacialis* | Northern fulmar | Aves | Procellariidae | Kerbiriou; Le Viol; Bonnet; Robert | Popul Ecol | 2012 | 10.1007/s10144-012-0306-9 | Colonial | 6.666 |
| *Giraffa camelopardalis* | Giraffe | Mammalia | Giraffidae | Lee; Bond; Kissui; Kiwango; Bolger | J Mammal | 2016 | 10.1093/jmammal/gyw086 | Social | 13.592 |
| *Gorgonia ventalina* | Common sea fan | Anthozoa | Gorgoniidae | Sabat; Toledo-Hernández | J Marin Biol | 2015 | 10.1155/2015/987060 | Colonial | -2.120 |
| *Gorilla beringei beringei* | Mountain gorilla | Mammalia | Hominidae | Morris; Altmann; Brockman; Cords; Fedigan; Pusey; Stoinski; Bronikowski; Alberts; Strier | Am Nat | 2011 | 10.1086/657443 | Social | 12.101 |
| *Gyps coprotheres* | Cape vulture | Aves | Accipitridae | Monadjem; Wolter; Neser; Kane | Anim Conserv | 2013 | 10.1111/acv.12054 | Communal | 8.970 |
| *Haliaeetus albicilla* | White-tailed eagle | Aves | Accipitridae | Krüger; Grünkorn; Struwe-Juhl | Biol Conserv | 2010 | 10.1016/j.biocon.2009.12.010 | Solitary | 8.475 |
| *Haliaeetus leucocephalus* | Bald eagle | Aves | Accipitridae | Etterson; Gerald Ankley | Environ Sci Technol | 2021 | 10.1021/acs.est.1c04791 | Solitary | 8.464 |
| *Halichoerus grypus* | Gray seal | Mammalia | Phocidae | Harwood | J Appl Ecol | 1978 | 10.2307/2402601 | Social | 12.194 |
| *Haliotis laevigata* | Greenlip abalone | Gastropoda | Haliotidae | Fordham; Mellin; Russell; Akçakaya; Bradshaw; Aiello-Lammens; Caley; Connell; Mayfield; Shepherd; Brook | Glob Change Biol | 2013 | 10.1111/gcb.12289 | Solitary | 3.822 |
| *Haliotis rufescens* | Red abalone | Gastropoda | Haliotidae | Rogers-Bennett; Leaf | Ecol Appl | 2006 | 10.1890/04-1688 | Solitary | 6.362 |
| *Homo sapiens* | Human | Mammalia | Hominidae | Keyfitz; Flieger | Book Chapter | 1990 | 0-226-43237-8 | Social | 11.084 |
| *Hoplocephalus bungaroides* | Broad-headed snake | Reptilia | Elapidae | Webb; Brook; Shine | Ecol Res | 2002 | 10.1046/j.1440-1703.2002.00463.x | Solitary | 5.631 |
| *Hybognathus argyritis* | Western Silvery Minnow | Actinopterygii | Cyprinidae | Young; Koops | Fish & Oceans Can | 2013e | NA | Gregarious | 6.024 |
| *Isurus oxyrinchus* | Shortfin mako shark | Elasmobranchii | Lamnidae | Tsai; Sun; Punt; Liu | ICES J Mar Sci | 2014 | 10.1093/icesjms/fsu056 | Solitary | 12.246 |
| *Kinosternon flavescens* | Yellow mud turtle | Reptilia | Kinosternidae | Iverson | Herpetologica | 1991 | 10.2307/1447431 | Solitary | 5.803 |
| *Kinosternon integrum* | Mud turtle | Reptilia | Kinosternidae | Macip-Ríos; Brauer-Robleda; Zúñiga-Vega; Casas-Andreu | Herpetol J | 2011 | NA | Solitary | 6.162 |
| *Kinosternon subrubrum* | Common mud turtle | Reptilia | Kinosternidae | Frazer; Gibbons; Greene | Ecology | 1991 | 10.2307/1941572 | Solitary | 5.041 |
| *Lagopus muta* | Japanese rock ptarmigan | Aves | Phasianidae | Suzuki; Kobayashi; Nakamura; Takasu | Wildlife Biol | 2013 | 10.2981/13-021 | Solitary | 6.256 |
| *Lagothrix lagotricha* | Humboldt's woolly monkey | Mammalia | Atelidae | Defler | Book Chapter | 2014 | 978-1-4939-0697-0 | Social | 8.743 |
| *Lepetodrilus fucensis* | Deep-sea limpet | Gastropoda | Lepetodrilidae | Kelly; Metaxas | Mar Ecol Prog Ser | 2010 | 10.3354/meps08442 | Communal | -2.120 |
| *Leptogorgia virgulata* | Colourful sea whip | Anthozoa | Gorgoniidae | Gotelli | Ecology | 1991 | 10.1046/j.1461-0248.2000.00138.x | Colonial | 7.496 |
| *Lepus europaeus* | European hare | Mammalia | Leporidae | Marboutin; Peroux | J Appl Ecol | 1995 | 10.2307/2404820 | Gregarious | 8.209 |
| *Lucanus miwai* | NA | Insecta | Lucanidae | Huang | J Asia-Pac Entomol | 2014 | 10.1016/j.aspen.2014.03.009 | Solitary | 4.382 |
| *Macaca mulatta* | Rhesus macaque | Mammalia | Cercopithecidae | Kessler; Pacheco; Rawlings; Ruiz-Lambrides; Delgado; Sabat | Am J Primatol | 2014 | 10.1002/ajp.22323 | Social | 8.493 |
| *Maccullochella peelii* | Murray cod | Actinopterygii | Percichthyidae | Stratford; Pollino; Brown | Environ Modell Softw | 2016 | 10.1016/j.envsoft.2016.02.009 | Solitary | 7.861 |
| *Macquaria ambigua* | Golden perch | Actinopterygii | Percichthyidae | Stratford; Pollino; Brown | Environ Modell Softw | 2016 | 10.1016/j.envsoft.2016.02.009 | Solitary | 8.248 |
| *Macrhybopsis storeriana* | Carmine shiner | Actinopterygii | Cyprinidae | Young; Koops | Fish & Oceans Can | 2013e | NA | Gregarious | 5.991 |
| *Marmota flaviventris* | Yellow-bellied marmot | Mammalia | Sciuridae | Ozgul; Oli; Armitage; Blumstein; Van Vuren | Am Nat | 2009 | 10.1086/597225 | Social | 8.219 |
| *Milvus migrans* | Black kite | Aves | Accipitridae | Sergio; Tavecchia; Blas; López; Tanferna; Hiraldo | Basic Appl Ecol | 2011 | 10.1016/j.baae.2010.11.004 | Colonial | 6.375 |
| *Mirounga angustirostris* | Northern elephant seal | Mammalia | Phocidae | Clinton; Le Boeuf | Ecology | 1993 | 10.2307/1939945 | Social | 13.726 |
| *Mirounga leonina* | Southern elephant seal | Mammalia | Phocidae | New; Clark; Costa; Fleishman; Hindell; Klanjšček; Lusseau; Kraus; McMahon; Robinson; Schick; Schwarz; Simmons; Thomas; Tyack; Harwood | Mar Ecol Prog Ser | 2014 | 10.3354/meps10547 | Social | 14.152 |
| *Moxostoma duquesnii* | Black redhorse | Actinopterygii | Catostomidae | Young; Koops | Fish & Oceans Can | 2014 | NA | Gregarious | 7.131 |
| *Mya arenaria* | Soft-shell clam | Bivalvia | Myidae | Brousseau | Mar Biol | 1978 | 10.1007/BF00390542 | Gregarious | 4.304 |
| *Mytilus californianus* | California mussel | Bivalvia | Mytilidae | Carson; Cook; Lopez-Duarte; Levin | Ecology | 2011 | 10.1890/11-0488.1 | Colonial | 2.303 |
| *Mytilus galloprovincialis* | Mediterranean mussel | Bivalvia | Mytilidae | Carson; Cook; Lopez-Duarte; Levin | Ecology | 2011 | 10.1890/11-0488.1 | Colonial | 2.079 |
| *Notropis anogenus* | Pugnose shiner | Actinopterygii | Cyprinidae | Venturelli; Vélez-Espino; Koops | Fish & Oceans Can | 2010 | NA | Gregarious | 1.099 |
| *Notropis photogenis* | Silver shiner | Actinopterygii | Cyprinidae | Young; Koops | Fish & Oceans Can | 2012a | NA | Gregarious | 3.219 |
| *Nuttallia obscurata* | Varnish clam | Bivalvia | Psammobiidae | Dudas; Dower; Anholt | Ecology | 2007 | 10.1890/06-1216.1 | Solitary | 2.251 |
| *Odocoileus virginianus* | White-tailed deer | Mammalia | Cervidae | Edmunds | Thesis | 2013 | NA | Solitary | 11.087 |
| *Oncorhynchus clarkii* | NA | Actinopterygii | Salmonidae | Murphy; Walsworth; Belmont; Conner; Budy | Ecosphere | 2020 | 10.1002/ecs2.3023 | Solitary | 7.741 |
| *Oncorhynchus tshawytscha* | Chinook salmon | Actinopterygii | Salmonidae | Fujiwara; Mohr; Greenberg | Acta Biotheo | 2014 | 10.1371/jourNAl.pone.0085464 | Gregarious | 8.434 |
| *Onychogalea fraenata* | Bridled nailtail wallaby | Mammalia | Macropodidae | Fisher; Hoyle; Blomberg | Ecol Appl | 2000 | 10.2307/2641054 | Solitary | 8.156 |
| *Opsopoeodus emiliae* | Pugnose minnow | Actinopterygii | Cyprinidae | Young; Koops | Fish & Oceans Can | 2012a | NA | Gregarious | 0.548 |
| *Orcinus orca* | Killer whale | Mammalia | Delphinidae | Vélez-Espino; Ford; Araújo; Ellis; Parken; Balcomb | Can Tech Report Fish & Aq Sci | 2014 | 978-1-100-23563-9 | Social | 15.274 |
| *Oreamnos americanus* | Mountain goat | Mammalia | Bovidae | Festa-Bianchet; Urquhart; Smith | Can J Zool | 1994 | 10.1139/z94-004 | Gregarious | 10.373 |
| *Otus scops* | Eurasian Scops Owl | Aves | Strigidae | Barbraud; Christian Barvoux; Guy Burneleau | Ibis | The demography of an increasing insular Eurasian Scops Owl (Otus scops) population in western France | 10.1111/ibi.12995 | Solitary | 4.905 |
| *Ovis aries* | Soay sheep | Mammalia | Bovidae | Clutton-Brock; Price; Albon; Jewell | J Anim Ecol | 1992 | 10.2307/5330 | Gregarious | 11.608 |
| *Ovis canadensis* | Bighorn sheep | Mammalia | Bovidae | Rubin; Boyce; Caswell-Chen | J Wildlife Manage | 2002 | 10.2307/3803144 | Gregarious | 11.144 |
| *Pan troglodytes schweinfurthii* | Eastern chimpanzee | Mammalia | Hominidae | Morris; Altmann; Brockman; Cords; Fedigan; Pusey; Stoinski; Bronikowski; Alberts; Strier | Am Nat | 2011 | 10.1086/657443 | Social | 10.597 |
| *Panthera pardus* | Leopard | Mammalia | Felidae | Balme; Slotow; Hunter | Biol Conserv | 2009 | 10.1016/j.biocon.2009.06.020 | Solitary | 10.779 |
| *Papio cynocephalus* | Olive baboon | Mammalia | Cercopithecidae | Morris; Altmann; Brockman; Cords; Fedigan; Pusey; Stoinski; Bronikowski; Alberts; Strier | Am Nat | 2011 | 10.1086/657443 | Social | 9.879 |
| *Paramuricea clavata* | Violescent sea-whip; Red gorgonian | Anthozoa | Plexauridae | Linares; Doak | Mar Ecol Prog Ser | 2010 | 10.3354/meps08437 | Colonial | 8.088 |
| *Pelagia noctiluca* | NA | Scyphozoa | Pelagiidae | Tomlinson; Maynou; Sabatés; Fuentes; Canepa; Sastre | Estuar Coast Shelf S | 2015 | 10.1016/j.ecss.2015.11.012 | Communal | 3.219 |
| *Percina copelandi* | Channel darter | Actinopterygii | Percidae | Venturelli; Vélez-Espino; Koops | Fish & Oceans Can | 2010 | NA | Gregarious | 1.099 |
| *Pernis apivorus* | European honey buzzard | Aves | Accipitridae | Bijlsma; Vermeulen; Hemerik; Klok | Ardea | 2012 | 10.5253/078.100.0208 | Solitary | 6.637 |
| *Petauroides volans* | Greater glider | Mammalia | Pseudocheiridae | Lindenmayer; Lacy; Pope | Ecol Appl | 2000 | 10.2307/2641117 | Solitary | 6.802 |
| *Phacochoerus aethiopicus* | Desert warthog | Mammalia | Suidae | Rodgers | Mammalia | 1984 | 10.1515/mamm.1984.48.3.327 | Solitary | 11.202 |
| *Nannopterum auritus* | Double-crested cormorant | Aves | Phalacrocoracidae | Chastant; King; Weseloh; Moore | J Wildlife Manage | 2014 | 10.1002/jwmg.628 | Colonial | 7.537 |
| *Phascolarctos cinereus* | Koala | Mammalia | Phascolarctidae | Rhodes; Ng; de Villiers; Preece; McAlpine; Possingham | Biol Conserv | 2011 | 10.1016/j.biocon.2010.12.027 | Solitary | 8.681 |
| *Phocarctos hookeri* | New Zealand sea lion | Mammalia | Otariidae | Meyer; Robertson; Chilvers; Krkošek | Mar Biol | 2015 | 10.1007/s00227-015-2695-8 | Social | 12.519 |
| *Phoebastria immutabilis* | Laysan albatross | Aves | Diomedeidae | Finkelstein; Doak; Nakagawa; Sievert; Klavitter | Anim Conserv | 2009 | 10.1111/j.1469-1795.2009.00311.x | Colonial | 7.833 |
| *Picoides borealis* | Red-cockaded woodpecker | Aves | Picidae | Maguire; Wilhere; Dong | J Wildlife Manage | 1995 | 10.2307/3802460 | Social | 3.871 |
| *Pimephales promelas* | Fathead minnow | Actinopterygii | Cyprinidae | Gleason | Hum Ecol Risk Assess | 2001 | 10.1080/20018091094835 | Colonial | 1.609 |
| *Podocnemis expansa* | Arrau turtle | Reptilia | Podocnemididae | Mogollones; Rodríguez; Hernández; Barreto | Chelonian Conserv Bi | 2010 | 10.2744/CCB-0778.1 | Solitary | 10.158 |
| *Poecilia reticulata* | Trinidadian guppy | Actinopterygii | Poeciliidae | Bronikowski; Clark; Rodd; Reznick | Ecology | 2002 | 10.1890/0012-9658(2002)083[2194:PDCOPI]2.0.CO;2 | Social | -0.693 |
| *Pongo abelii* | Sumatran orangutan | Mammalia | Hominidae | Wich; Utami-Atmoko; Setia; Rijksen; Schürmann; van Hooff; Van Schaik | J Hum Evol | 2004 | 10.1016/j.jhevol.2004.08.006 | Social | 10.886 |
| *Presbytis thomasi* | Thomas's langur | Mammalia | Cercopithecidae | Wich; Steenbeek; Sterck; Korstjens; Willems; Van Schaik | Am J Primatol | 2007 | 10.1002/ajp.20386 | Social | 8.808 |
| *Propithecus verreauxi* | Verreaux's sifaka | Mammalia | Indriidae | Morris; Altmann; Brockman; Cords; Fedigan; Pusey; Stoinski; Bronikowski; Alberts; Strier | Am Nat | 2011 | 10.1086/657443 | Social | 8.185 |
| *Puffinus auricularis* | Townsend's shearwater | Aves | Procellariidae | Martinez-Gomez; Jacobsen | Biol Conserv | 2004 | 10.1016/S0006-3207(03)00171-X | Colonial | 6.040 |
| *Puma concolor* | Cougar | Mammalia | Felidae | Lambert; Wielgus; Robinson; Katnik; Cruickshank; Clarke; Almack | J Wildlife Manage | 2006 | 10.2193/0022-541X(2006)70[246:CPDAVI]2.0.CO;2 | Solitary | 10.268 |
| *Pylodictis olivaris* | Flathead catfish | Actinopterygii | Ictaluridae | Sakaris; Irwin | Ecol Appl | 2010 | 10.1890/08-0305.1 | Solitary | 10.929 |
| *Rangifer tarandus* | Reindeer | Mammalia | Cervidae | Bjorkvoll; Lee; Grøtan; Saether; Stien; Engen; Albon; Loe; Hansen | Ecology | 2016 | 10.1890/15-0317.1 | Social | 11.180 |
| *Rutilus rutilus* | Common roach | Actinopterygii | Cyprinidae | Otjacques; De Laender; Kestemont | Ecol Model | 2016 | 10.1016/j.ecolmodel.2015.12.002 | Solitary | 7.518 |
| *Saguinus fuscicollis* | Saddlebacked tamarin | Mammalia | Callitrichidae | Watsa | Thesis | 2013 | 10.7936/K7DB7ZTD | Social | 5.976 |
| *Saguinus imperator* | Emperor tamarin | Mammalia | Callitrichidae | Watsa | Thesis | 2013 | 10.7936/K7DB7ZTD | Social | 6.178 |
| *Salvelinus confluentus* | Bull trout | Actinopterygii | Salmonidae | Bowerman | Thesis | 2013 | NA | Solitary | 8.040 |
| *Salvelinus malma* | Lake trout | Actinopterygii | Salmonidae | Spromberg; Birge | Environ Toxicol Chem | 2005 | 10.1897/04-160.1 | Gregarious | 9.815 |
| *Sardina pilchardus* | Sardine | Actinopterygii | Clupeidae | Serghini; Boutayeb; Auger; Charouki; Ramzi; Ettahiri; Tchente | Acta Biotheo | 2009 | 10.1007/s10441-009-9090-0 | Gregarious | 4.205 |
| *Scolytus ventralis* | Fir engraver beetle | Insecta | Curculionidae | Berryman | Can Entomol | 1973 | 10.4039/Ent1051465-11 | Solitary | 3.219 |
| *Sigmodon hispidus* | Hispid cotton rat | Mammalia | Muridae | Sauer; Slade | J Mammal | 1985 | 10.2307/1381244 | Solitary | 5.073 |
| *Somateria mollissima* | Common Eider | Aves | Anatidae | Öst; Ramula; Linden; Karell; Kilpi | Popul Ecol | 2016 | 10.1007/s10144-015-0517-y | Colonial | 7.646 |
| *Spermophilus dauricus* | Daurian ground squirrel | Mammalia | Sciuridae | Luo; Fox | J Mammal | 1990 | 10.2307/1381947 | Gregarious | 5.298 |
| *Sprattus sprattus* | European sprat | Actinopterygii | Clupeidae | Haslob; Hauss; Petereit; Clemmesen; Kraus; Peck | Mar Biol | 2012 | 10.1007/s00227-012-1933-6 | Gregarious | 3.684 |
| *Sterna hirundo* | Common tern | Aves | Laridae | Szostek | J Ornithol | 2011 | 10.1007/s10336-011-0745-7 | Colonial | 4.787 |
| *Sternotherus odoratus* | Common musk turtle | Reptilia | Kinosternidae | Mitchell | Herpetol Monogr | 1988 | 10.2307/1467026 | Solitary | 4.927 |
| *Sternula antillarum* | California least tern | Aves | Laridae | Massey; Bradley; Atwood | Condor | 1992 | 10.2307/1369293 | Colonial | 4.007 |
| *Suricata suricatta* | Meerkat | Mammalia | Herpestidae | Conquet; Ozgul; Blumstein; Armitage; Oli; Martin; Clutton-Brock; Paniw | Ecosphere | 2023 | 10.1002/ecy.3894 | Social | 6.654 |
| *Sus scrofa* | Wild boar | Mammalia | Suidae | Gamelon; Gaillard; Servanty; Gimenez; Toïgo; Baubet; Klein; Lebreton | J Appl Ecol | 2012 | 10.1111/j.1365-2664.2012.02160.x | Solitary | 11.813 |
| *Tamiasciurus hudsonicus* | American red squirrel | Mammalia | Sciuridae | McAdam; Boutin; Sykes; Humphries | Ecoscience | 2007 | 10.2980/1195-6860(2007)14[362:LHOFRS]2.0.CO;2 | Solitary | 5.513 |
| *Thalassarche melanophrys* | Black-browed albatross | Aves | Diomedeidae | Arnold; Brault; Croxall | Ecol Appl | 2006 | 10.1890/03-5340 | Colonial | 8.143 |
| *Thalia democratica* | NA | Thaliacea | Salpidae | Henschke; Smith; Everett; Suthers | J Planck Res | 2015 | 10.1093/plankt/fbv024 | Solitary | -2.166 |
| *Theropithecus gelada* | geladas | Mammalia | Cercopithecidae | Evan T Sloan; Jacinta C Beehner; Thore J Bergman; Amy Lu; Noah Snyder-Mackler; Jacquemyn | Ecol Evol | 2022 | 10.1002/ece3.8759 | Social | 9.678 |
| *Tympanuchus cupido* | Greater prairie chicken | Aves | Phasianidae | Fefferman; Reed | J Wildlife Manage | 2006 | 10.2307/3803419 | Solitary | 6.786 |
| *Umbonium costatum* | NA | Gastropoda | Trochidae | Noda; Nakao | J Anim Ecol | 1996 | 10.2307/5722 | Communal | 0.833 |
| *Urocitellus armatus* | Uinta ground squirrel | Mammalia | Sciuridae | Oli; Slade; Dobson | Ecology | 2001 | 10.1890/0012-9658(2001)082[1921:EODROU]2.0.CO;2 | Colonial | 5.738 |
| *Urocitellus beldingi* | Belding's ground squirrel | Mammalia | Sciuridae | Sherman; Morton | Ecology | 1984 | 10.2307/1939140 | Colonial | 5.667 |
| *Urocitellus columbianus* | Columbian ground squirrel | Mammalia | Sciuridae | Dobson; Oli | Am Nat | 2001 | 10.1086/321322 | Colonial | 6.150 |
| *Ursus americanus* | American black bear | Mammalia | Ursidae | Hebblewhite; Percy; Serrouya | Biol Conserv | 2003 | 10.1016/S0006-3207(02)00341-5 | Solitary | 11.794 |
| *Ursus arctos* | Brown bear | Mammalia | Ursidae | Wielgus | Biol Conserv | 2002 | 10.1016/S0006-3207(01)00265-8 | Solitary | 11.783 |
| *Ursus maritimus* | Polar bear | Mammalia | Ursidae | Hunter; Caswell; Runge; Regehr; Amstrup; Stirling | Ecology | 2010 | 10.1890/09-1641 | Solitary | 12.204 |
| *Vipera aspis* | Asp viper | Reptilia | Viperidae | Altwegg; Dummermuth; Anholt; Flatt | Oikos | 2005 | 10.1111/j.0030-1299.2001.13723.x | Solitary | 5.081 |
| *Vulpes vulpes* | Red fox | Mammalia | Canidae | Devenish-Nelson; Harris; Soulsbury; Richards; Stephens | Oikos | 2013 | 10.1111/j.1600-0706.2012.20706.x | Social | 7.976 |
| *Xenosaurus grandis* | Crevice-dwelling lizard | Reptilia | Xenosauridae | Zuniga-Vega; Valverde; Rojas-Gonzalez; Lemos-Espinal | Copeia | 2007 | 10.1643/0045-8511(2007)7[324:AOTPDO]2.0.CO;2 | Solitary | 3.161 |
| *Xenosaurus platyceps* | Flathead knob-scaled lizard | Reptilia | Xenosauridae | Rojas-Gonzalez; Jones; Zúñiga-Vega; Lemos-Espinal | Amphibia-Reptilia | 2008 | 10.1163/156853808784124992 | Solitary | 2.976 |
| *Xenosaurus agrenon* | Knob-scaled lizard | Reptilia | Xenosauridae | Zamora-Abrego; Chang; Zuniga-Vega; Nieto-Montes de Oca; Johnson | Herpetologica | 2010 | 10.1655/09-005.1 | Solitary | 8.161 |
| *Yoldia notabilis* | NA | Bivalvia | Yoldiidae | Nakaoka | Oikos | 1997 | 10.2307/3546090 | Colonial | 1.121 |
| *Zoarces viviparus* | European eelpout | Actinopterygii | Zoarcidae | Bergek; Ma; Vetemaa; Franzén; Appelberg | Ecotox Environ Safe | 2012 | 10.1016/j.ecoenv.2012.01.019 | Solitary | 6.234 |

**Table S2**. Loadings of the seven life history traits used to produce the phylogenetic PCA space described by Figure 5. The life history traits are: Generation time (*T*); Age at maturity (*L_α_*); Mean life expectancy (*η_e_*); Maturity probability (*p_R_*); Reproductive window (*L_α-ω_*); Degree of parity (*S*); and Net reproductive output (*R_0_*). The bottom two rows provided the eigenvalues associated with each principal component (PC) as well as the variance explained by each PCA.

|  | **PC1** | **PC2** | **PC3** | **PC4** | **PC5** | **PC6** | **PC7** |
| --- | --- | --- | --- | --- | --- | --- | --- |
| *T* | -0.19 | 0.813 | -0.193 | -0.259 | 0.338 | 0.291 | -0.013 |
| *L_α_* | -0.49 | 0.707 | -0.191 | 0.337 | -0.308 | -0.054 | -0.109 |
| *η_e_* | 0.583 | 0.712 | -0.262 | -0.045 | -0.183 | -0.143 | 0.172 |
| *p_R_* | 0.838 | -0.08 | 0.104 | -0.165 | -0.332 | 0.376 | -0.046 |
| *L_α-ω_* | 0.805 | 0.244 | -0.008 | -0.406 | 0.108 | -0.315 | -0.129 |
| S | 0.247 | -0.371 | -0.878 | 0.16 | 0.061 | 0.032 | -0.02 |
| *R_0_* | 0.662 | 0.184 | 0.251 | 0.648 | 0.207 | 0.031 | -0.011 |
| **Eigenvalue** | 1.981 | 1.316 | 0.966 | 0.801 | 0.503 | 0.404 | 0.069 |
| **% variance** | 33.7% | 26.2% | 16.6% | 12.5% | 5.7% | 4.6% | 0.8% |

**Table S3**. Scores of the phylogenetic PCA for each of the species in the study.

| Species | **PC1** | **PC2** | **PC3** | **PC4** | **PC5** | **PC6** | **PC7** |
| --- | --- | --- | --- | --- | --- | --- | --- |
| *Acipenser fulvescens* | -1.855 | -0.917 | 1.594 | -0.298 | 1.079 | 0.318 | -0.402 |
| *Acropora downingi* | -2.313 | -1.004 | -1.468 | 0.953 | -0.824 | 0.179 | 0.067 |
| *Ailuropoda melanoleuca* | 1.734 | -0.574 | -0.328 | 0.31 | -0.457 | -0.266 | -0.188 |
| *Alces alces* | 1.663 | -1.81 | -0.407 | -1.003 | -0.242 | -0.467 | 0.052 |
| *Alouatta seniculus* | 2.132 | 1.276 | 0.401 | -0.875 | -0.536 | -0.372 | -0.256 |
| *Ammocrypta pellucida* | 0.22 | -0.741 | -0.103 | 1.028 | 0.257 | -0.563 | -0.271 |
| *Anthropoides paradiseus* | 1.558 | 1.185 | 0.556 | 1.65 | 0.268 | 0.53 | 0.056 |
| *Astroblepus ubidiai* | -0.632 | -0.075 | 1.254 | 0.46 | -0.193 | -0.381 | -0.126 |
| *Bostrychia hagedash* | 0.73 | -0.308 | -0.675 | -0.805 | -0.635 | 0.165 | -0.031 |
| *Brachyrhaphis rhabdophora* | -0.78 | -0.449 | 0.601 | -1.766 | 1.43 | 0.209 | 0.161 |
| *Brachyteles hypoxanthus* | 0.648 | 2.475 | 0.408 | -0.329 | -0.73 | 0.85 | -0.256 |
| *Buteo solitarius* | -0.463 | 0.559 | -1.021 | -0.576 | 0.08 | 0.304 | 0.03 |
| *Callinectes sapidus* | -1.222 | -1.298 | 1.087 | -0.486 | -0.539 | -0.765 | -0.041 |
| *Callorhinus ursinus* | 0.71 | -0.434 | -0.498 | -0.835 | 0.129 | -0.289 | 0.057 |
| *Callospermophilus lateralis* | 2.676 | -2.577 | 0.935 | 0.247 | -0.803 | 0.564 | -0.168 |
| *Calyptorhynchus lathami* | 1.052 | 2.052 | -0.02 | -0.809 | -0.471 | -0.2 | -0.03 |
| *Canis lupus* | 2.415 | -1.35 | -0.038 | 0.582 | 0.137 | -0.286 | -0.024 |
| *Caretta caretta* | -2.474 | 0.755 | -0.028 | 0.23 | -0.211 | 0.66 | 0.663 |
| *Catostomus platyrhynchus* | -0.741 | -1.992 | -0.691 | -0.22 | 0.332 | -0.523 | -0.312 |
| *Cebus capucinus* | 0.957 | 3.031 | 0.52 | -1.013 | -0.382 | 0.451 | -0.27 |
| *Centrocercus minimus* | -1.016 | -1.155 | 0.663 | -0.512 | -0.466 | -0.471 | -0.039 |
| *Cephaloleia fenestrata* | -2.747 | 0.595 | 0.3 | 0.366 | -0.831 | 0.631 | -0.233 |
| *Cercopithecus mitis* | 1.582 | 3.057 | 0.192 | -0.734 | -0.943 | 0.372 | -0.319 |
| *Certhia americana* | 0.602 | -0.844 | -0.011 | -0.087 | -0.646 | -0.137 | -0.094 |
| *Cervus canadensis* | 0.984 | -1.559 | -0.868 | -0.744 | 0.609 | -0.496 | 0.139 |
| *Cervus elaphus* | 0.905 | -1.654 | -0.724 | -0.704 | 0.482 | -0.377 | 0.113 |
| *Chelonia mydas* | -2.274 | 0.274 | -0.689 | 0.99 | -0.151 | 0.725 | 0.038 |
| *Chelydra serpentina* | -2.738 | -0.038 | -1.7 | 0.918 | -0.179 | 1.02 | -0.725 |
| *Anser caerulescens* | 2.406 | -0.286 | -0.089 | -0.932 | -0.441 | 0.132 | -0.035 |
| *Chrysemys picta* | -1.056 | 0.295 | -2.017 | 0.317 | -0.871 | 0.256 | 0.15 |
| *Clemmys guttata* | -2.766 | 0.981 | -0.13 | 0.346 | -0.632 | 0.692 | -0.075 |
| *Myodes rufocanus* | 0.83 | -0.865 | 0.511 | 0.299 | -0.232 | -0.004 | -0.127 |
| *Colias alexandra* | -1.76 | 1.643 | 0.358 | 0.606 | -0.923 | 0.206 | 0.317 |
| *Coragyps atratus* | 0.672 | 2.661 | 0.832 | -0.15 | -0.714 | -0.139 | 0.075 |
| *Cottus aturi* | -0.63 | -1.612 | -0.595 | -0.14 | 0.482 | -0.361 | -0.311 |
| *Crocodylus acutus* | -2.07 | -0.712 | -1.173 | 1.032 | 0.387 | 0.869 | -0.149 |
| *Crocodylus johnsoni* | -0.57 | 0.976 | -0.521 | 0.476 | 1.021 | 0.441 | -0.277 |
| *Crocodylus niloticus* | -1.706 | 1.102 | 1.201 | 0.989 | 0.648 | 0.395 | -0.763 |
| *Cyprinus carpio* | 2.229 | -0.526 | 0.509 | 1.457 | 0.617 | 0.132 | -0.086 |
| *Didelphis aurita* | 2.463 | -0.99 | 1.591 | 1.351 | -0.185 | 0.788 | -0.353 |
| *Elephas maximus* | 3.353 | -1.745 | -0.295 | 1.264 | 0.749 | -0.377 | 0.218 |
| *Epidalea calamita* | -0.654 | -0.095 | -0.649 | 0.342 | 0.551 | 0.136 | -0.314 |
| *Erimyzon sucetta* | -0.486 | -2.22 | 2.059 | 0.678 | 0.785 | -0.82 | -0.203 |
| *Eumetopias jubatus* | -0.219 | -1.06 | -0.634 | -0.585 | -0.162 | -0.121 | 0.003 |
| *Falco peregrinus* | -0.031 | -0.501 | -0.472 | -1.009 | 0.565 | 0.117 | 0.083 |
| *Fulmarus glacialis* | 0.452 | 3.259 | -0.053 | -1.496 | -1.102 | -0.089 | -0.093 |
| *Giraffa camelopardalis* | -2.408 | -1.077 | 1.508 | -0.428 | 0.054 | 0.01 | -0.056 |
| *Gorgonia ventalina* | -1.078 | 0.66 | 0.514 | -0.98 | -0.064 | -0.182 | -0.029 |
| *Gorilla beringei beringei* | 1.154 | 2.113 | 0.914 | -0.07 | -0.791 | 0.568 | -0.501 |
| *Gyps coprotheres* | 1.831 | 1.324 | 0.325 | -0.911 | -0.28 | -0.507 | -0.18 |
| *Haliaeetus albicilla* | 0.519 | 0.385 | -1.692 | -0.619 | -0.57 | 0.214 | -0.107 |
| *Haliaeetus leucocephalus* | -0.575 | 0.854 | -1.072 | 0.156 | -0.432 | 0.359 | 0.162 |
| *Halichoerus grypus* | 2.222 | 1.287 | 0.184 | -0.349 | 0.248 | -0.724 | -0.437 |
| *Haliotis laevigata* | 0.496 | 0.554 | -1.602 | 0.169 | 0.117 | -0.203 | 0.052 |
| *Haliotis rufescens* | -1.212 | -0.015 | -0.092 | 0.123 | 0.184 | 0.166 | -0.128 |
| *Homo sapiens* | 3.937 | -0.11 | -0.152 | 0.476 | -0.581 | -0.186 | -0.139 |
| *Hoplocephalus bungaroides* | -0.674 | 0.72 | -0.979 | 0.677 | -0.269 | 0.358 | 0.069 |
| *Hybognathus argyritis* | 0.059 | -1.647 | 1.445 | 1.262 | 0.612 | -0.813 | -0.236 |
| *Isurus oxyrinchus* | -0.39 | 1.844 | 0.802 | 1.606 | -0.196 | 0.006 | 0.535 |
| *Kinosternon flavescens* | 1.145 | 1.655 | 1.37 | 1.012 | -0.012 | 1.887 | 0.139 |
| *Kinosternon integrum* | -2.001 | 0.043 | -1.848 | 0.191 | 0.257 | 0.902 | -0.314 |
| *Kinosternon subrubrum* | -1.036 | 0.163 | -2.211 | 0.277 | 0.213 | 0.597 | -0.212 |
| *Lagopus muta* | 2.481 | -2.195 | 0.221 | -1.005 | -0.992 | 0.424 | 0.141 |
| *Lagothrix lagotricha* | 0.444 | 1.706 | -1.917 | -0.232 | -0.262 | 0.067 | -0.09 |
| *Lepetodrilus fucensis* | -0.511 | -0.124 | 0.55 | -1.479 | 0.596 | -0.248 | 0.258 |
| *Leptogorgia virgulata* | -1.181 | -0.623 | 1.506 | -2.163 | 2.045 | 0.492 | 0.165 |
| *Lepus europaeus* | -0.924 | -1.961 | 1.287 | -0.501 | 0.077 | -0.716 | -0.1 |
| *Lucanus miwai* | -0.053 | 1.576 | 1.223 | 1.773 | -0.436 | 0.059 | 0.26 |
| *Macaca mulatta* | 3.672 | 2.14 | 0.802 | 0.158 | 0.126 | -0.247 | -0.482 |
| *Maccullochella peelii* | 0.704 | 0.595 | -0.237 | 2.317 | 1.014 | 0.389 | -0.132 |
| *Macquaria ambigua* | 1.079 | -0.11 | 1.38 | 1.48 | 0.624 | -0.122 | -0.017 |
| *Macrhybopsis storeriana* | 1.684 | -3.164 | -0.202 | -0.802 | -0.734 | 1.117 | -0.56 |
| *Marmota flaviventris* | 0.671 | -1.618 | -0.367 | -0.737 | -0.346 | 0.201 | -0.102 |
| *Milvus migrans* | 2.481 | -1.624 | -0.538 | -1.255 | -0.283 | 1.042 | 0.04 |
| *Mirounga angustirostris* | -0.992 | -1.074 | -0.847 | 0.109 | -0.298 | 0.271 | -0.007 |
| *Mirounga leonina* | -0.575 | -0.049 | 0.842 | -0.462 | -0.522 | -0.202 | 0.215 |
| *Moxostoma duquesnii* | -2.138 | 0.203 | 1.013 | -1.132 | 1.031 | 0.789 | -0.257 |
| *Mya arenaria* | 1.871 | 0.109 | -0.364 | 1.166 | 1.009 | -0.456 | 0.154 |
| *Mytilus californianus* | -0.491 | -2.042 | -1.507 | 0.019 | -0.175 | -0.781 | -0.321 |
| *Mytilus galloprovincialis* | -0.383 | -1.784 | -1.163 | 0.202 | -0.033 | -0.72 | -0.314 |
| *Notropis anogenus* | -0.238 | -1.36 | -0.558 | 0.216 | -0.216 | -0.689 | -0.235 |
| *Notropis photogenis* | -1.238 | -0.202 | 0.729 | -0.004 | 0.178 | -0.104 | -0.315 |
| *Nuttallia obscurata* | -0.092 | -0.286 | -2.042 | -0.89 | 0.551 | 0.557 | -0.207 |
| *Odocoileus virginianus* | 0.806 | -1.918 | -0.251 | -0.365 | 0.153 | 0.409 | -0.087 |
| *Oncorhynchus clarkii* | -1.075 | -1.168 | 1.262 | 0.055 | -0.215 | -0.71 | -0.041 |
| *Oncorhynchus tshawytscha* | -1.617 | -1.494 | 1.502 | 0.185 | -0.231 | -0.595 | -0.29 |
| *Onychogalea fraenata* | 1.086 | 0.092 | -0.114 | 0.878 | -0.052 | -0.25 | -0.006 |
| *Opsopoeodus emiliae* | 1.637 | -1.993 | 1.502 | -0.887 | -1.705 | 0.63 | -0.268 |
| *Orcinus orca* | -0.431 | 1.226 | 0.195 | -0.912 | -0.064 | -0.195 | -0.193 |
| *Oreamnos americanus* | -1.016 | -0.783 | 0.847 | -0.389 | -1.228 | -0.2 | 0.126 |
| *Otus scops* | 3.005 | -0.722 | 0.07 | -1.153 | -0.22 | 0.795 | 0.132 |
| *Ovis aries* | 1.212 | -0.43 | 0.481 | -0.352 | 0.009 | -0.388 | 0.049 |
| *Ovis canadensis* | -0.168 | -0.744 | 0.146 | -1.056 | -0.039 | 0.015 | 0.067 |
| *Pan troglodytes schweinfurthii* | 0.877 | 0.551 | 0.643 | -1.986 | 0.09 | -0.285 | -0.326 |
| *Panthera pardus* | -0.375 | -0.73 | -0.564 | -0.044 | 0.145 | -0.03 | -0.021 |
| *Papio cynocephalus* | 1.387 | 2.357 | 0.924 | -0.169 | -0.148 | 0.199 | -0.393 |
| *Paramuricea clavata* | -1.429 | 0.759 | -1.78 | 0.362 | -0.975 | 0.26 | 0.455 |
| *Pelagia noctiluca* | -0.996 | -1.407 | 0.653 | -0.571 | -0.324 | -0.893 | -0.119 |
| *Percina copelandi* | 0.422 | -0.14 | 1.808 | 1.638 | 0.789 | -0.5 | -0.207 |
| *Pernis apivorus* | 0.607 | 1.026 | 0.497 | -1.185 | -1.023 | 0.02 | 0.092 |
| *Petauroides volans* | 0.157 | -0.452 | -1.362 | -0.685 | -0.442 | -1.577 | 0.619 |
| *Phacochoerus aethiopicus* | -0.665 | -1.855 | -0.646 | -0.318 | 0.198 | -0.237 | -0.143 |
| *Nannopterum auritus* | -0.496 | -0.116 | -0.542 | -0.073 | -0.005 | 0.087 | -0.013 |
| *Phascolarctos cinereus* | 3.717 | 0.962 | 0.43 | -0.972 | 0.06 | -0.331 | 0.023 |
| *Phocarctos hookeri* | 0.07 | -1.319 | -1.155 | -1.18 | 1.266 | 0.027 | 0.096 |
| *Phoebastria immutabilis* | -1.116 | 1.683 | -0.973 | 0.169 | -0.492 | 0.686 | 0.37 |
| *Picoides borealis* | 0.637 | -0.6 | -0.61 | -0.61 | -0.25 | 0.048 | 0.022 |
| *Pimephales promelas* | -0.216 | -0.819 | 0.343 | 0.558 | 0.379 | -0.517 | -0.242 |
| *Podocnemis expansa* | -1.835 | 0.497 | 0.863 | -1.101 | 0.971 | 0.465 | 0.056 |
| *Poecilia reticulata* | -0.61 | 0.105 | 0.062 | -1.694 | 1.287 | 0.299 | 0.149 |
| *Pongo abelii* | 1.562 | 0.336 | -1.922 | -0.875 | -0.804 | -0.079 | -0.083 |
| *Presbytis thomasi* | 0.699 | 0.914 | -1.506 | -0.997 | -0.252 | -0.64 | -0.072 |
| *Propithecus verreauxi* | 1.463 | 2.885 | 0.266 | -2.021 | -0.809 | -0.51 | -0.653 |
| *Puffinus auricularis* | 0.536 | 1.876 | -0.209 | -0.924 | -0.483 | -0.041 | 0.063 |
| *Puma concolor* | 1.99 | -2.86 | -0.043 | -1.397 | -0.484 | 0.821 | -0.013 |
| *Pylodictis olivaris* | -2.834 | -0.207 | 1.264 | 0.474 | 0.406 | 0.967 | -0.351 |
| *Rangifer tarandus* | 3.502 | -0.297 | -0.88 | -0.684 | 0.092 | -0.421 | 0.132 |
| *Rutilus rutilus* | -1.486 | -1.185 | -0.144 | 0.463 | 0.207 | 0.003 | -0.476 |
| *Saguinus fuscicollis* | 2.171 | -0.408 | -0.411 | -0.312 | -0.353 | -0.038 | 0.028 |
| *Saguinus imperator* | 0.738 | -0.225 | -1.462 | -1.075 | 0.381 | 0.88 | 0.122 |
| *Salvelinus confluentus* | -1.877 | -0.826 | 0.094 | 0.352 | -0.212 | 0.014 | -0.244 |
| *Salvelinus malma* | -1.221 | -1.767 | 1.486 | 0.307 | 0.189 | -0.596 | -0.278 |
| *Sardina pilchardus* | -0.747 | -1.26 | 1.723 | -0.312 | -0.504 | -0.948 | 0.015 |
| *Scolytus ventralis* | 0.011 | 1.769 | 0.762 | 1.803 | -0.105 | -0.14 | 0.271 |
| *Sigmodon hispidus* | -0.791 | -1.737 | 1.403 | -0.039 | -0.214 | -1.067 | -0.087 |
| *Somateria mollissima* | 2.411 | 1.415 | -0.727 | -3.104 | 0.08 | 0.492 | -0.251 |
| *Spermophilus dauricus* | 0.192 | -1.354 | -0.266 | -0.435 | -0.248 | -0.135 | -0.097 |
| *Sprattus sprattus* | 0.871 | 3.419 | -2.172 | -1.162 | 0.404 | -1.269 | -0.373 |
| *Sterna hirundo* | 4.234 | 0.363 | 0.307 | -0.042 | 0.856 | 0.987 | 0.081 |
| *Sternotherus odoratus* | -2.292 | 0.156 | -0.078 | -0.356 | 0.431 | 0.947 | -0.201 |
| *Sternula antillarum* | 1.077 | 0.267 | -1.377 | -0.228 | -0.342 | -0.021 | -0.011 |
| *Suricata suricatta* | 0.102 | 0.224 | 0.698 | -1.402 | -0.833 | 0.634 | -0.004 |
| *Sus scrofa* | 2.515 | -0.492 | -0.769 | 1.612 | 0.821 | -0.989 | -0.196 |
| *Tamiasciurus hudsonicus* | 2.047 | -2.291 | 0.053 | -1.05 | -0.83 | 0.884 | -0.1 |
| *Thalassarche melanophrys* | -2.527 | 1.058 | -1.84 | 0.425 | -0.064 | 1.448 | 0.335 |
| *Thalia democratica* | 4.7 | 1.266 | -0.664 | -0.488 | -0.148 | 0.334 | 0.127 |
| *Theropithecus gelada* | 1.708 | 2.248 | 0.642 | -1.542 | -0.739 | -0.619 | -0.414 |
| *Tympanuchus cupido* | 0.412 | -1.438 | 1.447 | 0.176 | -0.29 | -0.445 | -0.016 |
| *Umbonium costatum* | 1.922 | 0.516 | -0.88 | -0.172 | -0.396 | 0.294 | 0.015 |
| *Urocitellus armatus* | 1.842 | -2.838 | 1.178 | -0.855 | -1.398 | 0.427 | -0.104 |
| *Urocitellus beldingi* | 0.725 | -1.75 | -0.939 | 0.086 | -0.205 | -0.086 | -0.203 |
| *Urocitellus columbianus* | -0.139 | -0.558 | 0.569 | -0.606 | -0.893 | -0.175 | 0.044 |
| *Ursus americanus* | -0.789 | -0.114 | 0.494 | -0.495 | -0.32 | -0.082 | 0.177 |
| *Ursus arctos* | 1.037 | 0.224 | -1.397 | -0.261 | -0.498 | -0.255 | -0.22 |
| *Ursus maritimus* | -0.541 | 1.289 | -0.009 | -0.796 | -0.783 | -0.062 | 0.216 |
| *Vipera aspis* | -1.331 | 0.242 | -0.882 | -0.008 | -0.284 | 0.259 | -0.049 |
| *Vulpes vulpes* | 2.212 | -2.949 | 0.955 | -0.634 | -1.027 | 0.394 | -0.01 |
| *Xenosaurus grandis* | 1.029 | 0.381 | -0.78 | 0.518 | -0.6 | 0.148 | -0.083 |
| *Xenosaurus platyceps* | -0.306 | 0.475 | -0.774 | 0.313 | -0.336 | -0.005 | -0.004 |
| *Xenosaurus agrenon* | -1.833 | -1.292 | 1.656 | -0.6 | -0.137 | -0.518 | -0.211 |
| *Yoldia notabilis* | -0.23 | -0.139 | 1.521 | -0.842 | 1.351 | -0.059 | 0.211 |
| *Zoarces viviparus* | -0.667 | -1.633 | -0.82 | 0.779 | 0.392 | -0.259 | -0.499 |

**Figure S1**. Histogram of frequency of the 152 studies used in this study, from the COMADRE Animal Matrix Database, as a function of the frequency with which they were sampled within a year (Projection Interval, [[43]](https://paperpile.com/c/tNUvkn/d0cKJ)). To ensure comparability of the outputs across all species’ matrix population models (MPMs), each MPM was rescaled so time units were all on an annual basis (See Methods).

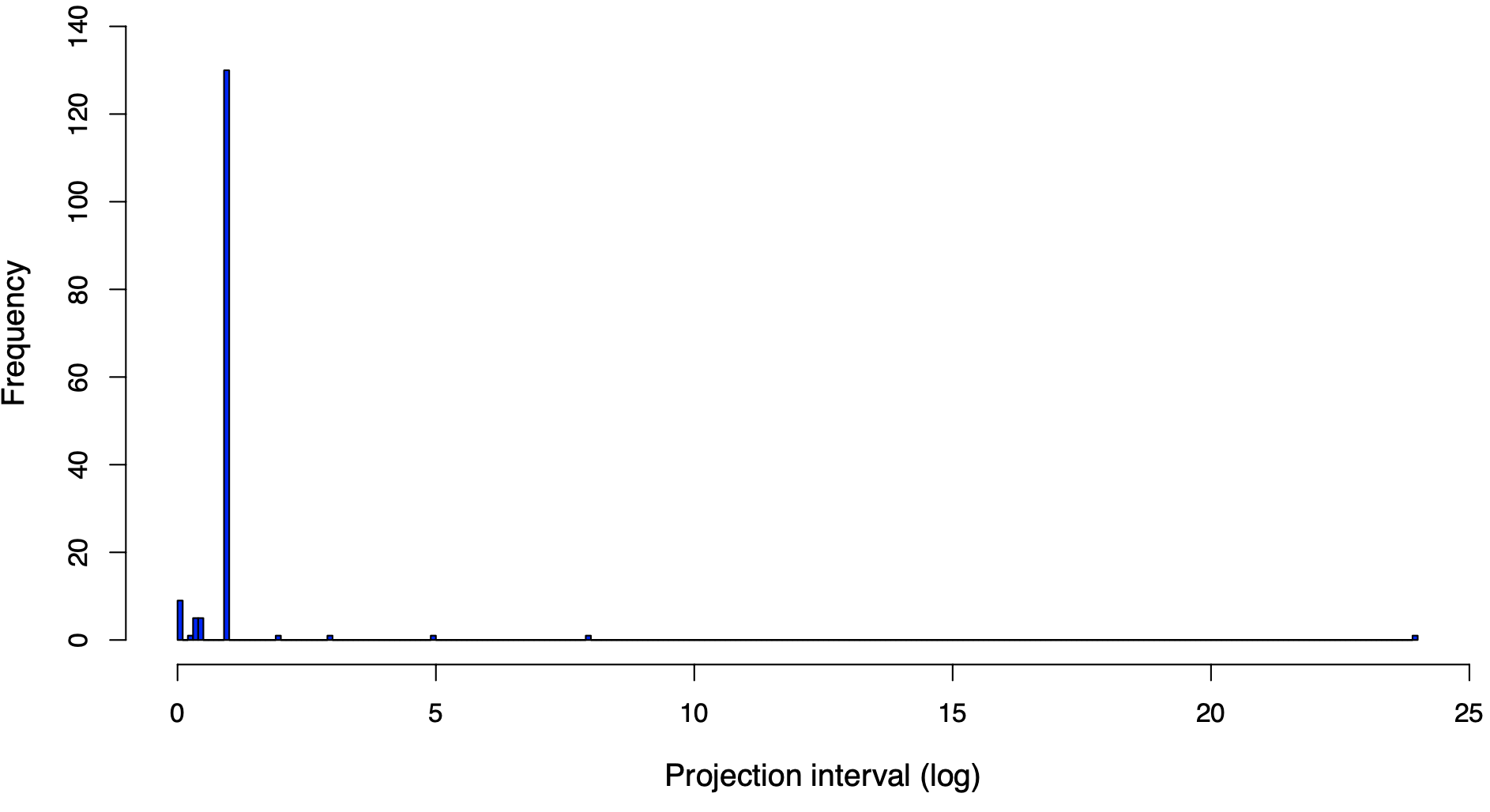

**Figure S2**. An exploration of the values of degree of parity (S) that emerge from different age-specific survivorship (*l_x_*) and reproductive (*m_x_*) schedules. The metric S ranges from 0 (for strictly semelparous schedules, at the left) to >>0 (for highly iteroparous schedules, at the right), regardless of the shape of the survivorship curve (type I, II, and III from top to bottom, respectively).

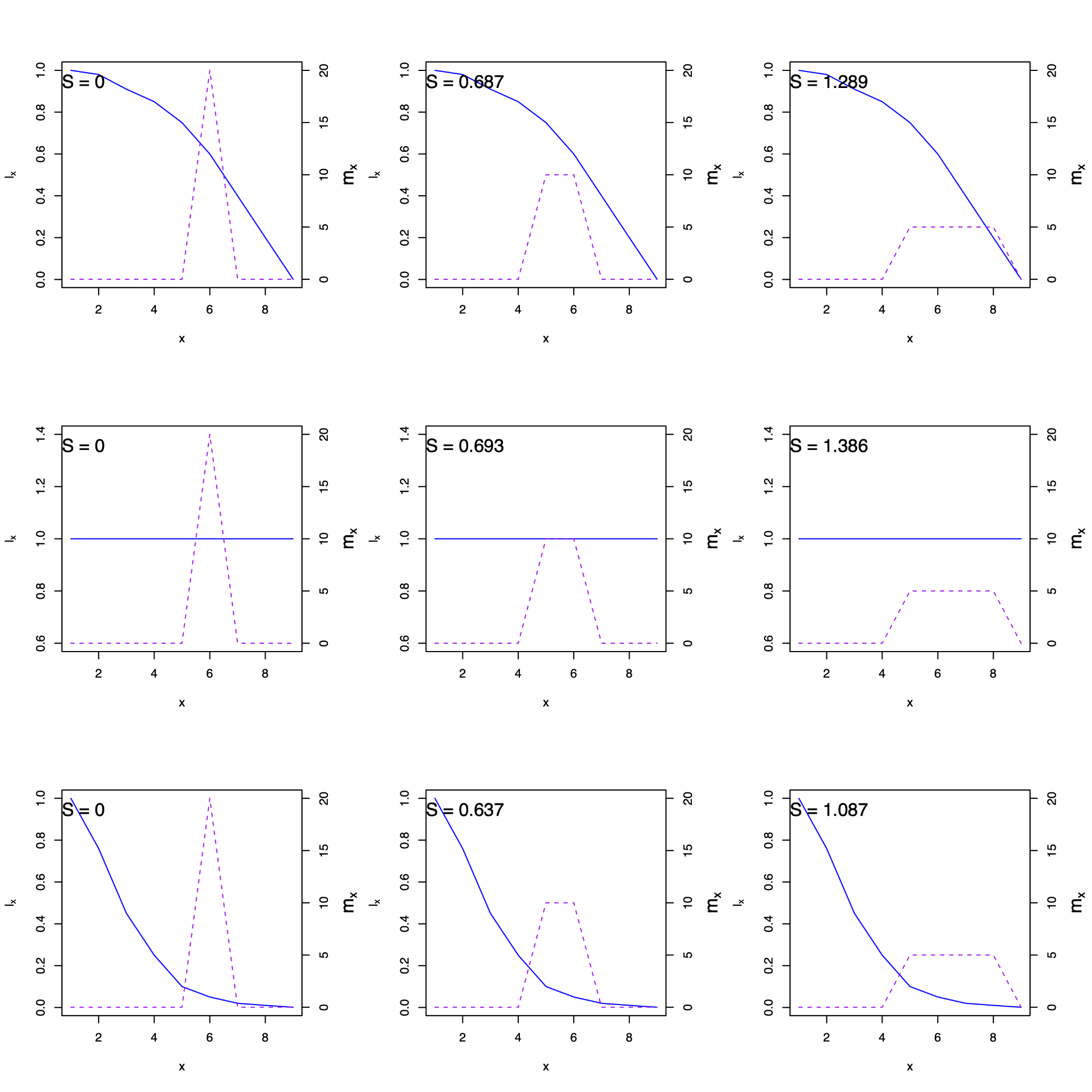

**Figure S3**. Pairwise correlation of the eleven life history traits considered in this study before phylogenetic imputations. The life history traits are: Generation time (*T*); Mean life expectancy (*η_e_*); Maximum longevity (*L_max_*); Variance in life expectancy (*Δη_e_*); Maturity probability (*p_R_*); Age at maturity (*L_α_*); Reproductive window (*L_α-ω_*); Net reproductive output (*R_0_*); Degree of parity (*S*); Reproductive senescence (*s_mx_*); and Actuarial senescence (*s_lx_*).

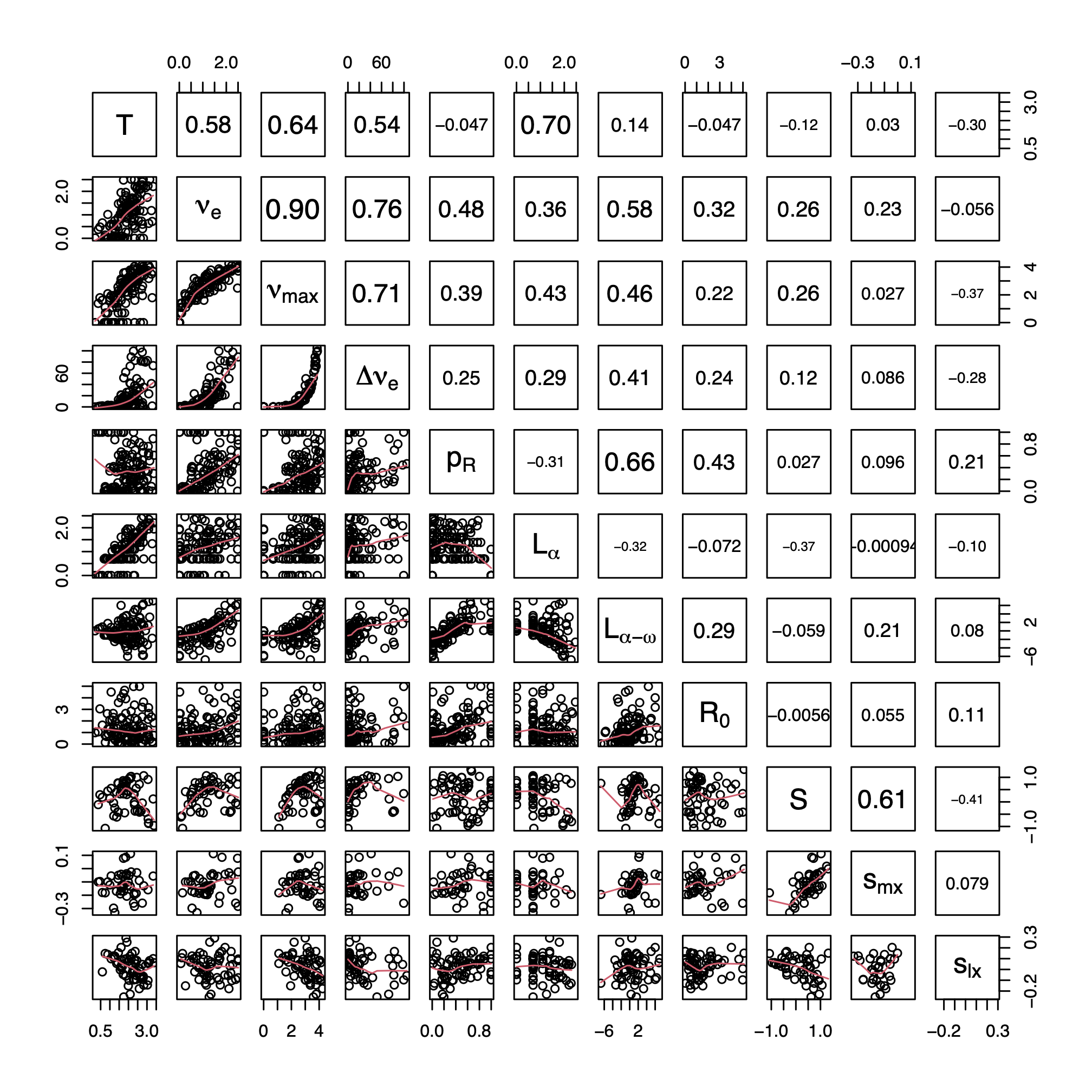

**Figure S4**. Percentage of missing data in each of the eight life history traits examined in the 152 animal species. Description of each life history trait symbol is provided in Table 1 and Fig. S3.

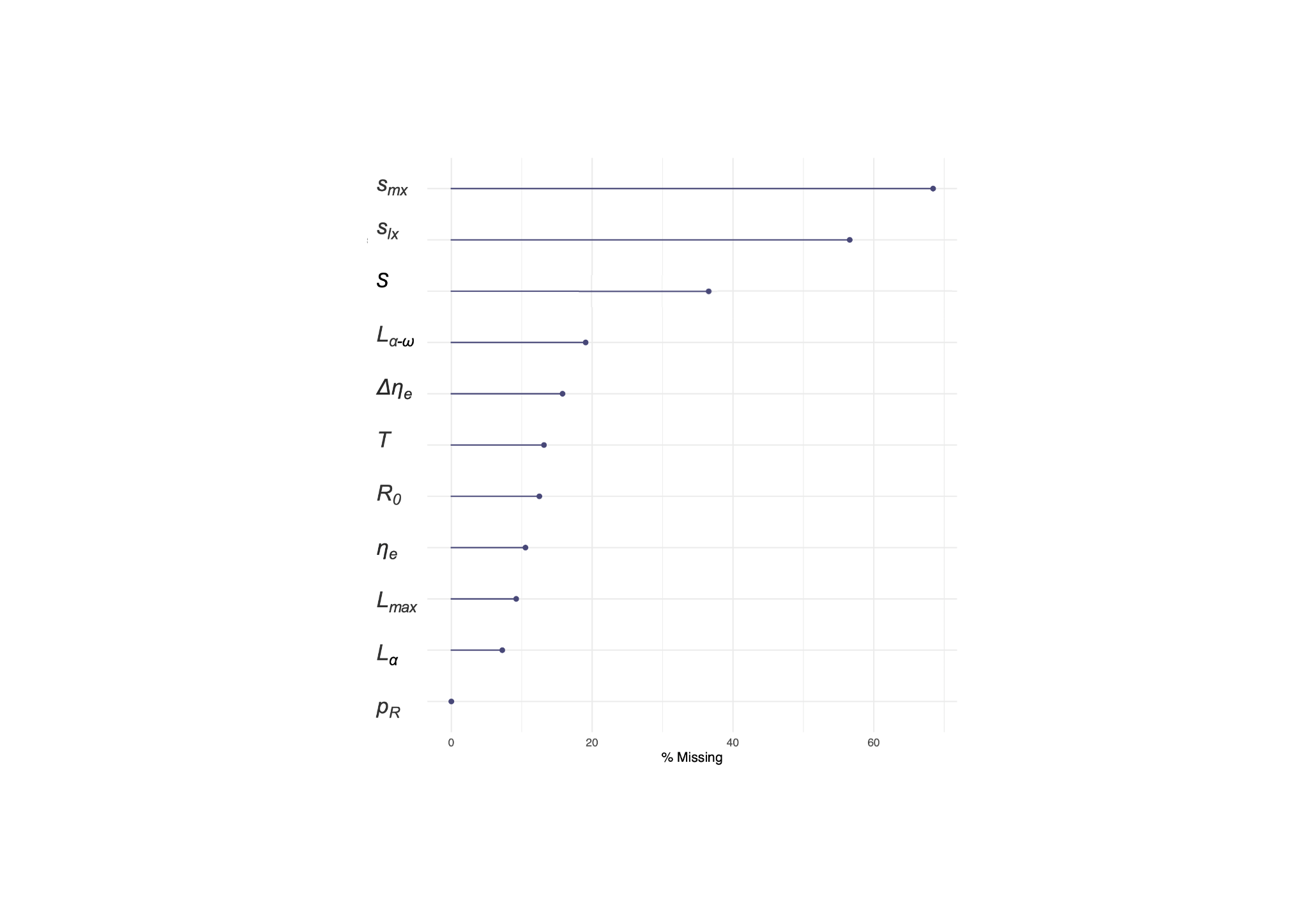

**Figure S5**. Pairwise correlation of the eleven life history traits considered in this study after phylogenetic imputations. The life history traits are defined in Fig. S3.

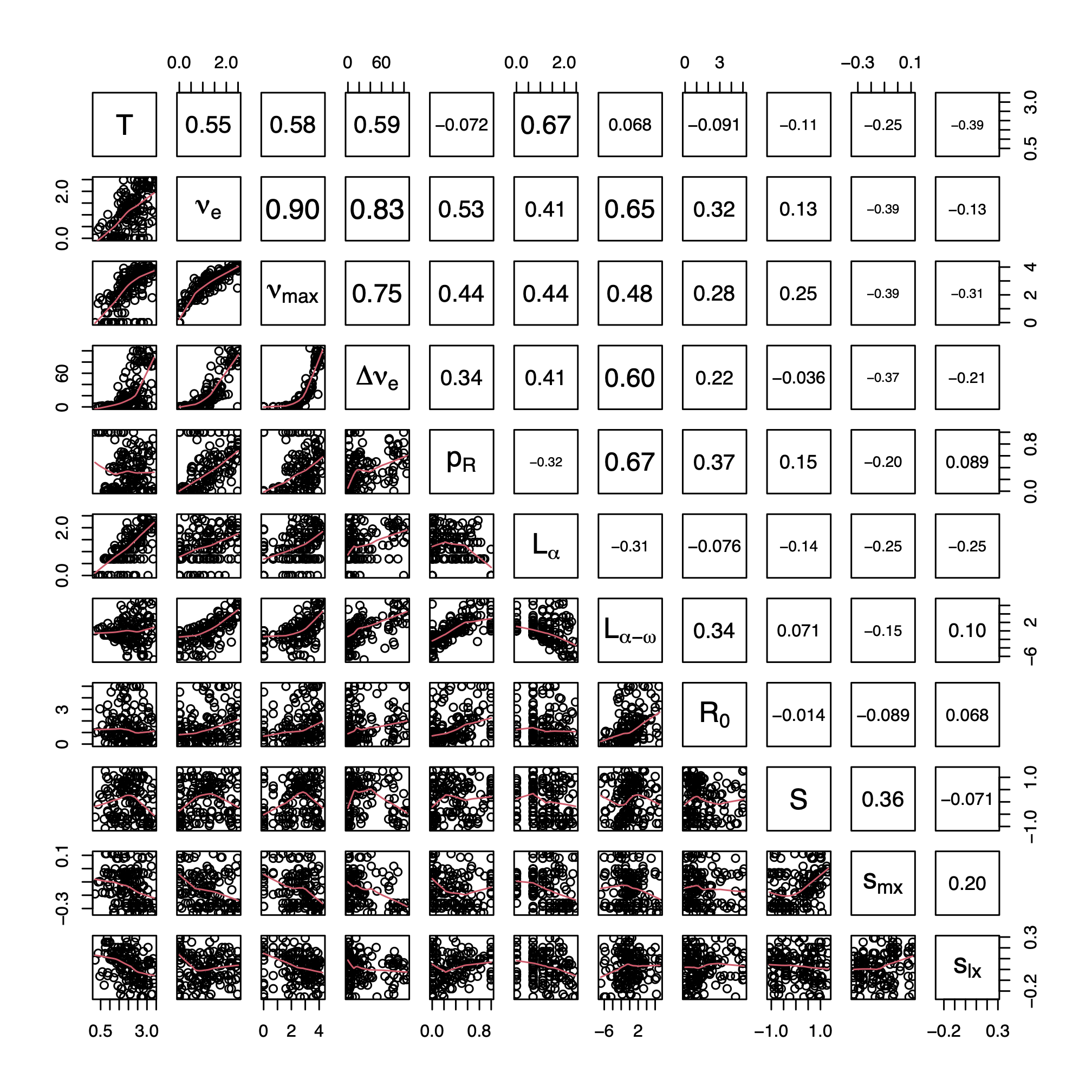

Additional Information

**Ethics**

NA.

**Data Accessibility**

The datasets supporting this article have been uploaded as part of the Supplementary Material. The rest of the data is found open-access at [www.compadre-db.org](http://www.compadre-db.org).

**Authors' Contributions**

RSG developed the ideas, digitised/obtained the data, performed the analyses, and wrote the manuscript.

**Competing Interests**

*I have no competing interests, other than the fact that I myself am a guest editor in this special feature*.
